# Phosphoproteomics of cellular mechanosensing reveals NFATC4 as a regulator of myofibroblast activity

**DOI:** 10.1101/2023.02.13.528335

**Authors:** Laura F. Mattner, Zhen Zeng, Christoph H. Mayr, Meshal Ansari, Xin Wei, Sara Asgharpour, Anita A. Wasik, Nikolaus Kneidinger, Mircea-Gabriel Stoleriu, Jürgen Behr, Julien Polleux, Ali Önder Yildirim, Gerald Burgstaller, Matthias Mann, Herbert B. Schiller

## Abstract

Feedback connections between tissue stiffness and cellular contractile forces can instruct cell identity and activity via a process referred to as mechanosensing. Specific phosphoproteome changes during mechanosensing are poorly characterized. In this work, we chart the global phosphoproteome dynamics of primary human lung fibroblasts sensing the stiffness of injury relevant fibronectin coated Poly(dimethylsiloxane) substrates. We discovered a key signaling threshold at a Young’s modulus of eight kPa stiffness, above which cells activated a large number of pathways including RhoA, CK2A1, PKA, AMPK, AKT1, and Hippo-YAP1/TAZ mediated signaling. Time-resolved phosphoproteomics of cell spreading on stiff substrates revealed the temporal dynamics of these stiffness-sensitive signaling pathways. ECM substrate stiffness above eight kPA induced fibroblast contractility, cytoskeletal rearrangements, ECM secretion, and a fibroblast to myofibroblast transition. Our data indicate that phosphorylation of the transcriptional regulator NFATC4 at S213/S217 enhances myofibroblast activity, which is the key hallmark of fibrotic diseases. NFATC4 knock down cells display reduced stiffness induced collagen secretion, cell contractility, nuclear deformation and invasion, suggesting NFATC4 as a novel target for antifibrotic therapy.

**Synopsis:** How tissue stiffness regulates identity and activity of tissue fibroblasts is unclear. Mass spectrometry based analysis of tissue stiffness dependent phosphoproteome changes reveals how primary lung fibroblasts sense the mechanical properties of their environment and identifies NFATC4 as a novel regulator of the stiffness dependent transition of fibroblasts to ECM secreting myofibroblasts.

- Mass spectrometry analysis reveals the signaling landscape of fibroblast mechanosensing
- Time-resolved phosphoproteomic analysis of cell spreading on fibronectin
- NFATC4 regulates myofibroblast collagen secretion, cell contractility and invasion

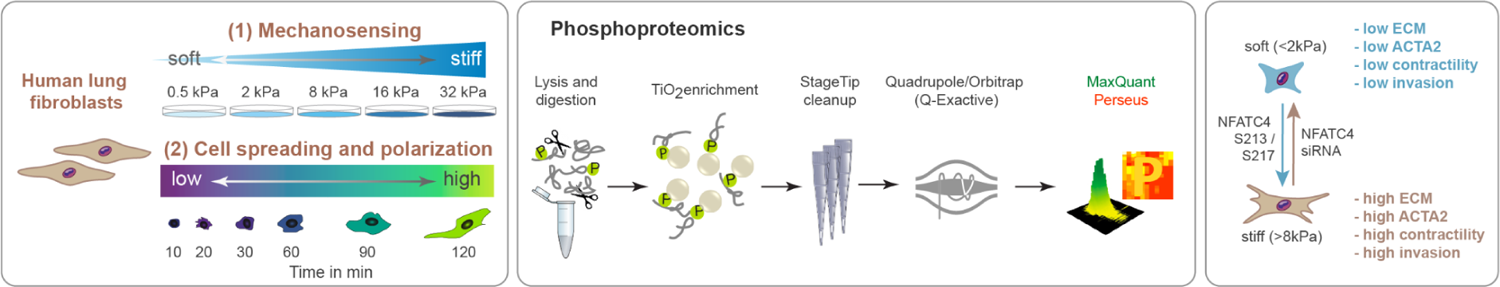

## Introduction

Cells can actively sense and respond to a variety of mechanical inputs from their environment (Vogel & Sheetz, 2006; Jansen *et al*, 2015). Tension and forces applied to cells are converted into chemical signals that enable cells to adopt their phenotype according to their surroundings (Jansen *et al*, 2015). Indeed, cellular responses to mechanical forces are substantial in embryonic development and adult physiology, and are involved in many diseases, such as atherosclerosis, hypertension, osteoporosis, muscular dystrophy, myopathies, cancer and metastasis formation (Hytönen & Wehrle-Haller, 2016; Hoffman *et al*, 2011). The molecular features that translate mechanical forces into biochemical signals are largely unknown and mass spectrometry driven proteomics can be used to reveal mechanosensitive proteome alterations in an unbiased manner (Wasik & Schiller, 2017).

Mechanosensitive signaling pathways are highly relevant in the pathophysiology of fibrotic diseases, where the composition of the extracellular matrix (ECM) and its mechanical properties are altered during disease progression (Zhou *et al*, 2013). A physiological stiffness range (0.2 – 2 kPa) keeps human lung fibroblasts in a quiescent state, while substrates with higher stiffness (2-35 kPa), as observed in fibrotic lungs, induce a profibrogenic phenotype with high proliferation and matrix synthesis rates (Liu *et al*, 2010). Thus, the number, activity and survival of activated fibroblasts in fibrotic diseases is at least to some extent controlled by mechanical signals. Cells encountering their extracellular matrix (ECM) substrate display ‘mechanoreciprocit’, which functions via feedback connections between cell-matrix adhesions and the cytoskeleton, adjusting the strength of the contractile forces to a balance of applied force and tensile strength of the ECM substrate (Schiller & Fässler, 2013).

The precise molecular nature of many elements in these mechanoreciprocal feedback connections is currently unknown. In most cases, the conversion of extracellular and mechanical stimuli into intracellular cues is encoded by post-translational modifications (PTMs). One of the five most common PTMs is protein phosphorylation (Walsh *et al*, 2005), which is a key modification for cellular signaling and is involved in nearly all biological functions (Needham *et al*, 2019). Importantly, protein phosphorylation on serine, threonine and tyrosine residues (S, T, Y) was shown to regulate protein conformation, as well as protein interactions and subcellular localization (Zaidel-Bar & Geiger, 2010), likely affecting proteins with functions in cell-matrix adhesions and the cytoskeleton.

Both integrin- and cadherin-mediated adhesions attach to the filamentous (F-) actin cytoskeleton using a diversity of adaptor and signaling proteins. These proteins join together to form a dense, highly dynamic network that is visible as a protein plaque on the plasma membrane, whose composition we call an adhesome. Remarkably, the recruitment of multiple plaque proteins into the adhesome requires myosin II-mediated mechanical tension (Kuo *et al*, 2011; Schiller *et al*, 2011). Thousands of phosphorylation sites are regulated in an integrin dependent manner (Schiller *et al*, 2013), and it is currently unclear which phosphorylation sites are directly regulated by mechanical forces.

In addition, cell shape plays a major role in cell fate decision and tissue morphogenesis (Nisenholz *et al*, 2014). The acquisition of cytoskeletal structure and polarity is organized by elastic stresses which develop in the cell during spreading onto their substrates (Zemel *et al*, 2011). Migration and cell spreading are controlled both by biochemical activity within the cell and by the stiffness of the underlying substrate and these activities are generally intensified on stiffer substrates (Schwarz & Safran, 2013). Through an increase in tension in the cytoskeleton, adhesion receptors are activated and the nucleus deformes, which can affect differentiation and growth control (Nisenholz *et al*, 2014).

In this work we ask how the stiffness of the ECM environment affects signaling and cell identity of human lung fibroblasts. We explore proteome-wide molecular changes in protein phosphorylation during mechanosensing and cell spreading using mass spectrometry driven phosphoproteomics (Humphrey *et al*, 2018). We discover a major switch in cellular signaling events at a stiffness of >8kPa that impacts a large number of kinases and signaling pathways. Integrative analysis of this data with time-resolved cell spreading data reveals the temporal dynamics of this cellular mechanosensing process. Finally, loss of function experiments show that the stiffness induced phosphorylation of the transcriptional regulator NFATC4 controls aspects of (myo-)fibroblast activity, which may have relevance in fibrotic disease.

## Results

### Matrix stiffness activates human lung fibroblasts

Fibroblasts from a normal healthy male human lung (CCL-151 cells) can be activated by matrix stiffness into a pro-fibrogenic state, characterized by increased proliferation and matrix synthesis rates (Liu *et al*, 2010). We seeded CCL-151 cells for 120 min on fibronectin coated PDMS substrates with a stiffness of 0.5 kPa and 28 kPa respectively and assessed the nuclear versus cytoplasmic localization of the mechanosensitive transcriptional regulator YAP1 (Figure 1A). YAP1 is known to translocate to the nucleus in a mechanical force and cell shape dependent manner (Elosegui-Artola *et al*, 2017; Nardone *et al*, 2017). Indeed, in CCL-151 cells we found nuclear shuttling on stiff 28 kPa PDMS substrates, while on soft 0.5 kPa substrates YAP1 was localized throughout the whole cell (Figure 1A).

**Figure 1.**
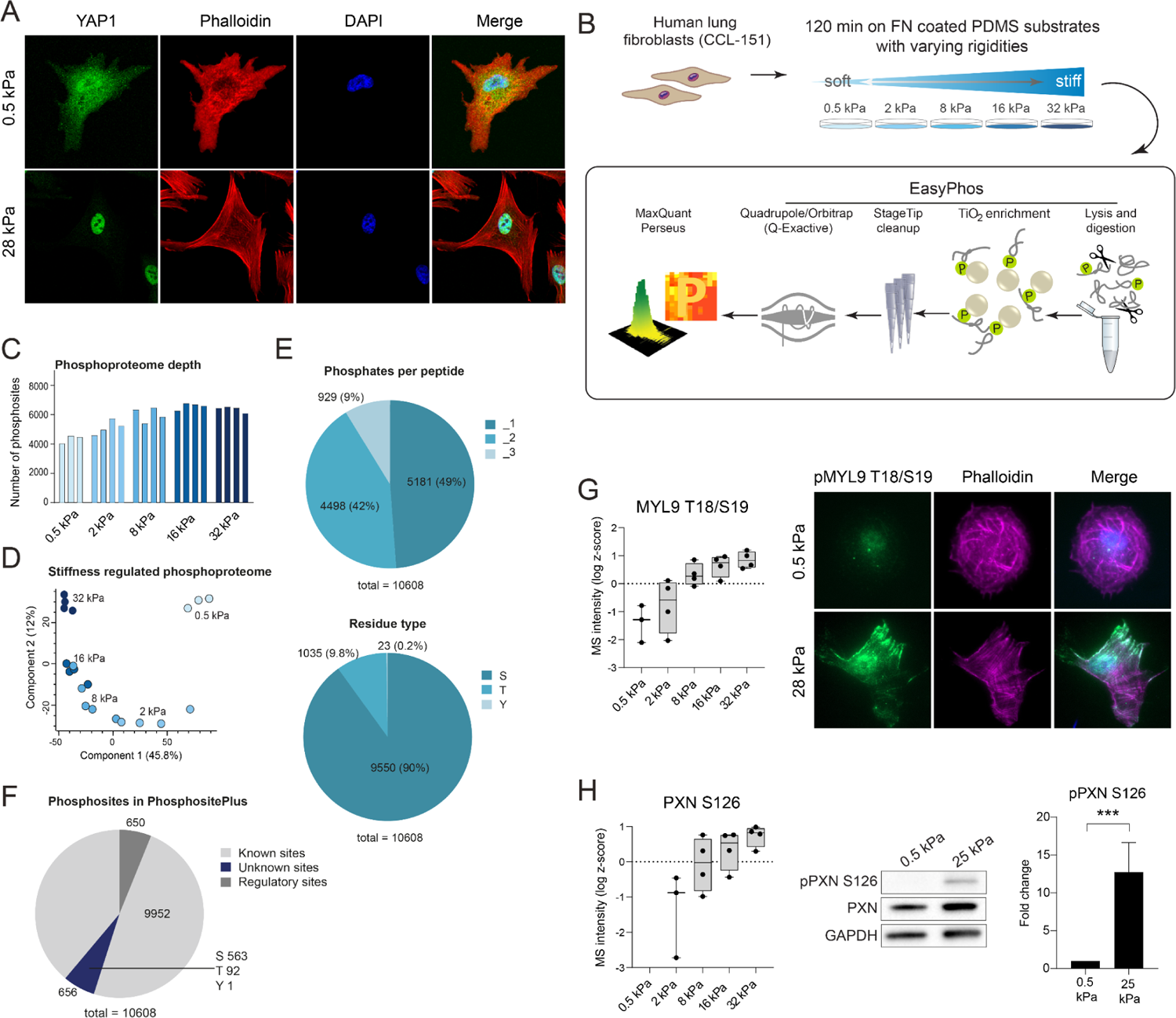
Human lung fibroblasts activate mechanosensitive signaling pathways on stiff ECM substrates. A. Immunofluorescence staining of YAP1 (green) in CCL-151 cells seeded for 2 h on fibronectin coated substrates with the indicated stiffness. B. Experimental workflow to analyze the phosphoproteome of human lung fibroblasts seeded on varying substrate stiffnesses. C. The bar graph shows the number of quantified phosphosites in the indicated experimental conditions. D. Principal component analysis (PCA) of the stiffness regulated phosphoproteome. E. Upper panel: the pie chart represents the percentage of identified phosphorylated serines (S), threonines (T) and tyrosines (Y) residues. Lower panel: the pie chart represents the percentage of identified singly, doubly and triply (or more) phosphorylated peptides. F. The pie chart shows the number of known sites, previously unreported sites and known sites with an assigned regulatory function in the PhosphoSitePlus database (www.phosphosite.org). G. The box plot shows phosphopeptide intensities of MYL9 T18/S19 at the indicated experimental conditions (box shows the range between the median of the lower half of the dataset and the upper half of the dataset and whiskers show minimum/maximum to the lower/upper quartile, respectively). Immunofluorescence staining with phosphospecific MYL9 T18/S19 antibody (green) confirms the stiffness dependent induction of phosphorylation. Phalloidin stain (magenta) shows the actin cytoskeleton, and nuclei (DAPI) are colored in blue. H. The box plot shows phosphopeptide intensities of PXN S126 at the indicated experimental conditions (box shows the and whiskers show minimum/maximum to the lower/upper quartile). Immunoblot analysis using a phosphospecific PXN S126 antibody and a PXN antibody confirms the stiffness induced induction of phosphorylation. GAPDH served as a loading control and was used to normalize the quantification (n=3, type of test - p-value).

We next generated deep phospho-proteomes of CCL-151 lung fibroblasts during stiffness sensing. CCL-151 cells were seeded for 120 min on fibronectin coated PDMS gels with varying rigidities from soft to stiff (0.5, 2, 8, 16 and 32 kPa) (Figure 1B). We identified 10,608 class-I phosphosites with a localization probability of the phosphosite > 0.75. In each replicate, included in the analysis, at least 4000 phosphosites were measured (Table S1). With increasing stiffness, more phosphorylation sites were recorded, in the range between 5400 and 6700 sites (Figure 1C). Principal component analysis of the phospho-peptide quantification showed a good separation of the distinct substrate rigidities and high reproducibility between the single replicates (Figure 1D). The 10,608 class-I phosphosites correspond to mainly serine residues (S) with 9550 sites, followed by 1035 phosphothreonines (T) and only 23 phosphorylated tyrosine residues (Y) (Figure 1E). Almost half of all identified phosphosites (5181) are localized on phosphopeptides which have only one phosphate group as modification. But 4498 phosphosites are found on doubly phosphorylated peptides and 929 sites are located on phosphopeptides with at least three phosphorylations (Figure 1E). We found a total of 656 phosphosites which were previously unknown without entry in the PhosphoSitePlus database. Further, 650 phosphorylation sites identified were already well characterized and known to be functionally relevant (Figure 1F). After filtering for sites at least measured three times in one of the stiffness conditions a total of 8349 phosphorylation sites remained and were used for further analysis.

To validate the experimental setup, we inspected well known mechanosensitive phosphosites. Phosphorylation of T19 and S20 on myosin regulatory light chain (MYL9) is needed for cells to increase their contractility (Schiller *et al*, 2013). Due to mechanoreciprocity, contractility of cells should increase on stiff substrates. In agreement with this prediction we observed a gradual increase in MYL9 T19/S20 with substrate stiffness in the phosphoproteome data, which we also validated using immunostainings (Figure 1G). Phosphorylation of serine 126 on the focal adhesion protein paxillin is known to regulate cytoskeletal remodeling and turnover of paxillin in focal adhesions (Cai *et al*, 2006). Consistent with the observed change in contractility and cytoskeletal architecture of CCL-151 on stiff substrates we also observed highly significant changes in PXN S126 in both phosphoproteomes and western blotting (Figure 1H).

In summary, the pro-fibrogenic phenotype of CCL-151 lung fibroblasts on stiff substrates (Liu *et al*, 2010) is accompanied by a massive remodeling of the phosphoproteome, including induction of the Yap signaling pathway, increased cell contractility and cytoskeletal rearrangements.

### The signaling landscape of fibroblast mechanosensing

To dive into the analysis of stiffness regulated signaling pathways, kinases and phosphatases, we performed an ANOVA test, which revealed 1631 significantly regulated phosphosites across the different substrate rigidities (1 % FDR) (Figure 2A, Table S2). These regulated sites account for around 20% of the phosphorylation sites in the entire data set and belong to 721 individual proteins. Mainly phosphosites with phosphorylation on serine residues were found to be regulated, namely 1463 sites, compared to 164 threonines and four tyrosines. More than half of the regulated sites increased phosphorylation on stiff substrates. We identified ten clusters containing between 52 to 648 phosphorylation sites, which showed different patterns of regulation (Figure 2A).

**Figure 2.**
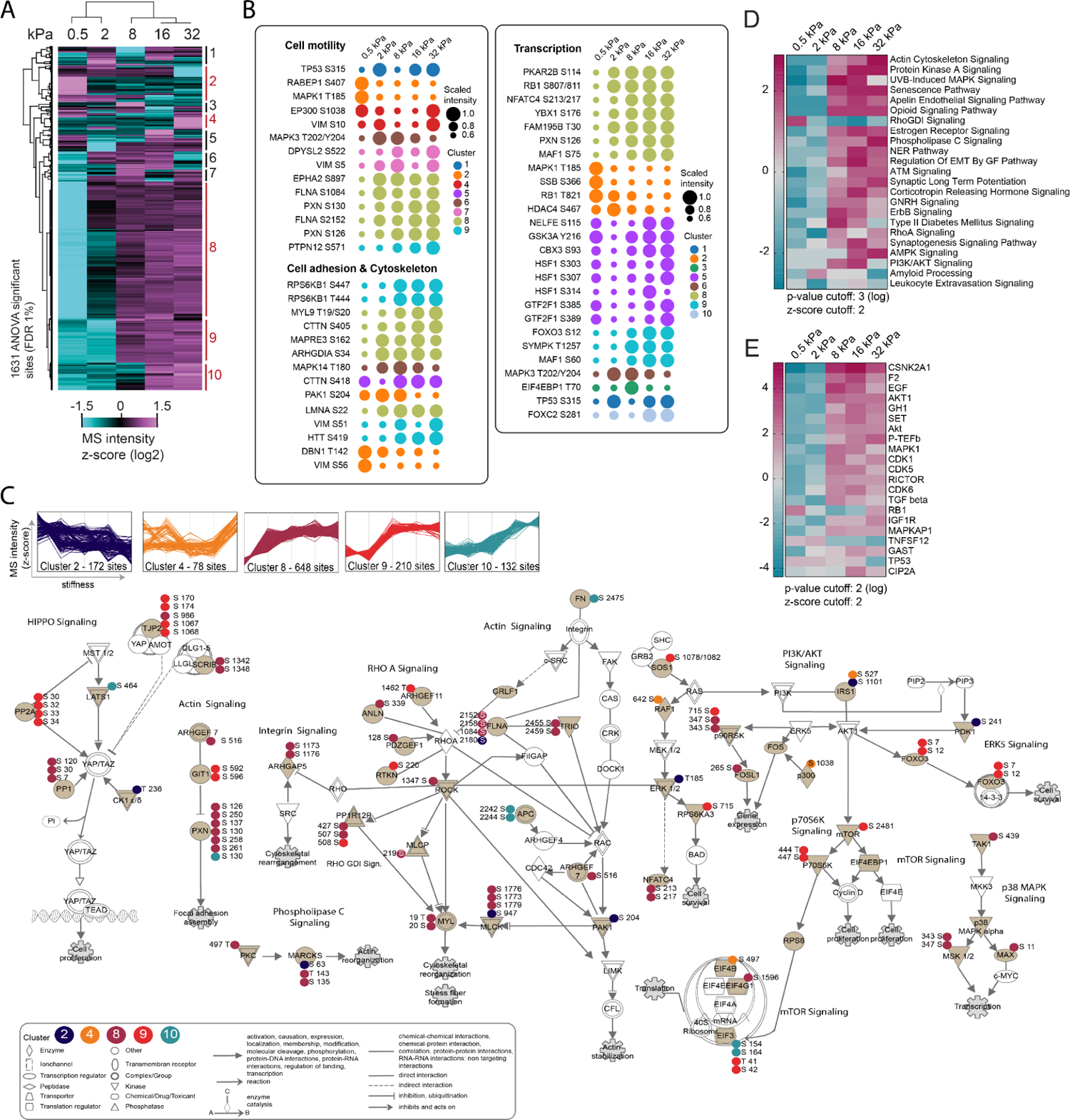
Phosphoproteomics reveals mechanosensitive signaling pathways. A. Heatmap depicts the difference of phosphorylation between the stiffness conditions by hierarchical clustering (Pearson correlation) of the z-scored MS intensities. B. Dot plots show the dynamics of scaled phosphorylation intensities on soft to stiff substrates for selected phosphosites with known regulatory site processes (PhosphositePlus database) associated with the indicated functional categories. Cluster identities from (A) are indicated by colors. C. The line plots show the dynamic regulation of the indicated clusters by substrate rigidity. The map shows functional relationships of a selected subset of stiffness regulated phosphosites in the indicated canonical signaling networks. The phosphorylation sites are color-coded by cluster ID. Nodes with shading in light brown represent proteins with at least one significantly regulated phosphosite. White shading of nodes represents proteins with no detected or not significantly regulated phosphosite. D–E. Phosphoproteomic data was analysed using Ingenuity Pathway Analysis (IPA). Significance of term enrichment is indicated through shading as log p-value. Regulated pathways belonging to “Canonical pathways” (D) and predicted “Upstream regulator” (E) with significant enrichment are shown.

Using the PhosphoSitePlus database (www.phosphosite.org) we identified 112 significantly regulated phosphosites with known regulatory function. This allowed us to assign stiffness regulated phosphosites to previously characterized molecular processes in cell adhesion, cytoskeletal reorganization, cell motility and regulation of transcription (Figure 2B). Several kinases involved in cell adhesion and cell motility were differentially phosphorylated on known functionally relevant sites. For instance, PAK1 S204, which regulates conformation and enzymatic activity of PAK1 and plays an important role in the formation of nascent adhesions and cell protrusions (Parrini *et al*, 2009), was downregulated on stiff substrates. Also the phosphorylation of ERK2 T185 was downregulated on stiff substrates, indicating higher ERK2 activity in physiologically soft environments (Singh *et al*, 2012). The PTPN12 S571 site, which regulates interaction of this phosphatase with the focal adhesion kinase (FAK) and thereby enables dephosphorylation of FAK Y397 (Zheng *et al*, 2011), was increased with stiffness. Also many cytoskeleton associated proteins were differentially phosphorylated at important regulatory sites. For instance, DBN1 T142 blocks the cryptic F-actin-bundling activity of the actin-filament binding protein Drebrin (Worth *et al*, 2013), which we found gradually decreased with increasing substrate stiffness (Figure 2B). This observation could partially explain the increased F-actin bundling and resulting stress fiber formation on stiff substrates (Figure 1G).

Alterations in ECM substrate stiffness could induce long-term gene expression changes via differential activity of transcription and chromatin remodeling factors. Sites with known regulatory functions on the transcriptional regulators RB1, PKAR2B, YBX1, NFATC4, FAM95B, MAF1, SYMPK and FOXO3a were gradually increased with substrate stiffness (Figure 2B). NFATC4 S213/S217 leads to the transactivation of the protein and increases transcriptional activity (Yao *et al*, 2007), indicating that this transcription factor gets activated by matrix stiffness in lung fibroblasts. SYMPK T1257 is a possible ERK target site and was increased on stiff substrates (Figure 2B). The protein is localized to tight junctions and the nucleus and is involved in transcriptional modulation and mRNA maturation as well as tight junction assembly. Interestingly, phosphorylation of SYMPK T1257 promotes nuclear localization of symplekin and interaction with YBX3, which enhances cell proliferation and differentiation (Zhang *et al*, 2017). Hyperphosphorylation of FOXO3 has been implicated with myofibroblasts in idiopathic pulmonary fibrosis (Al-Tamari *et al*, 2018). Consistent with this, we observed increased phosphorylation of FOXO3 S12 on stiff substrates.

Most significantly regulated phosphosites do not have known regulatory functions yet. To reveal possible connections between sites we mapped selected sites onto canonical signaling pathways (Figure 2C). Sections of the pathways HIPPO signaling, actin cytoskeleton signaling, integrin signaling, phospholipase C signaling, RHO A signaling, PI3K/AKT signaling, p70S6K signaling, ERK signaling, p38 MAPK signaling and mTOR signaling were manually merged and enriched with the phosphosite data from the rigidity sensing experiment. Selected sites were color coded according to the five most striking dynamic profiles, which consisted of sites that were generally higher on soft substrates and then clusters that started to emerge at different levels of rigidity (Figure 2C). Cluster 8 was the largest of the ten clusters and already showed a rise in phosphorylation at 2 kPa, whereas the phosphorylation in cluster 9 was only increased from a rigidity of 8 kPa. In contrast, the phosphosites contained in cluster 10 started to go up at a substrate stiffness of 16 kPa. The 78 phosphorylation sites present in cluster 4 only increased at 32 kPa. Sites in cluster 2 were highest on soft 0.5 kPa.

Finally, we searched for significantly altered pathways based on the phosphosite quantification to score potential upstream regulators in each of the rigidity conditions, using the Ingenuity Pathway Analysis platform (IPA, Qiagen, Redwood City, www.qiagen.com/ingenuity). The analyses predicted altered pathways (Figure 2D) and key upstream kinases (Figure 2E), and indicated that a stiffness of 8 kPa represents an important signaling threshold in lung fibroblasts at which the activity of kinases and associated regulation of signaling pathways was drastically altered. To browse the data we visualized predicted upstream kinases for selected mechanosensitive phosphosites (Figure S1). In an alternative approach, we used Metascape (Zhou *et al*, 2019) for pathway enrichment analysis (Figure S2A). The clusters 2, 8, 9 and 10 were enriched with sites corresponding to ‘supramolecular fiber organization’, ‘actin filament based process’ and ‘actin cytoskeleton organization’. Interestingly only cluster 2 (downregulated on stiff) contained phosphosites enriched for the term ‘focal adhesion’ and ‘fibroblast migration’. Cluster 10, which contains 132 sites upregulated on 16 and 32 kPa was significantly enriched for the terms ‘hippo signaling’, ‘mesenchymal cell differentiation’ and ‘Rho A signaling’. In contrast, the sites in cluster 8 (upregulated >2kPa) were enriched for annotations like ‘mRNA processing’, ‘regulation of mRNA stability’ and ‘RNA splicing’.

The casein kinase II (CK-II, CK2A1, CSNK2A1) was identified as the top upstream regulator in the IPA analysis (Figure 2E). This result was inferred based on the phosphorylation status of known CK2A1 targets (Figure S2B). A Fischer’s exact test was performed with the sites predicted by IPA to be CK2A1 targets to identify statistically enriched protein annotations (Figure S2C). Terms such as ‘regulation of translation’ and ‘mRNA processing’ and also the CK2A1 substrate motif were found to be overrepresented. To validate the *in silico* finding of enhanced CK-II activity *in vitro*, cell lysates of CCL-151 cells, which were seeded for 120 min on 0.5 kPa and 25 kPa substrates, were probed with an antibody specific for phosphorylated pS/pTDXE motif, a CK2A1 consensus sequence (Meggio & Pinna, 2003). This assay independently also predicted the stiffness dependent increase in CK2A1 activity, which was sensitive to the specific CK-II inhibitor TBB (Figure S2D).

To validate some of our observations in freshly isolated primary human lung fibroblasts (pHLF), we isolated pHLF from non-involved peritumor human lung tissue and repeated the phosphoproteome analysis in the same way as done for CCL-151 cells (Figure 1). Similar to CCL-151 the pHLF were spreading more on stiff substrates (Figure S3A). In pHLF we detected 7191 class-I phosphosites with a localization probability > 0.75. Principal component analysis showed a clear and reproducible separation of the distinct substrate rigidities (Figure S3B). The depth of pLFH phosphoproteomes was lower as in CCL-151 with between 2000 and 4000 quantified sites per replicate (Figure S3C-F). After filtering for sites that were at least measured four times in one of the stiffness conditions, 4725 sites remained for further analysis. Similar to CCL-151 (Figure 1) we found increased phosphorylation of MYL9 T10/S20 (Figure S3G) and PXN S126 (Figure S3H) when pHLF were cultured on stiff substrates. After filtering for significant sites, we identified 290 shared mechanosensitive phosphorylation sites in CCL-151 and pHLF (ANOVA significant with FDR <1%). In order to visualize the overlap in the kinetics across both experiments we embedded the PCA space of phosphosite intensities in one integrated UMAP plot. Using the louvain clustering algorithm we determined 7 distinct clusters of phosphosites in this integrated dataset (Figure 3A).

**Figure 3.**
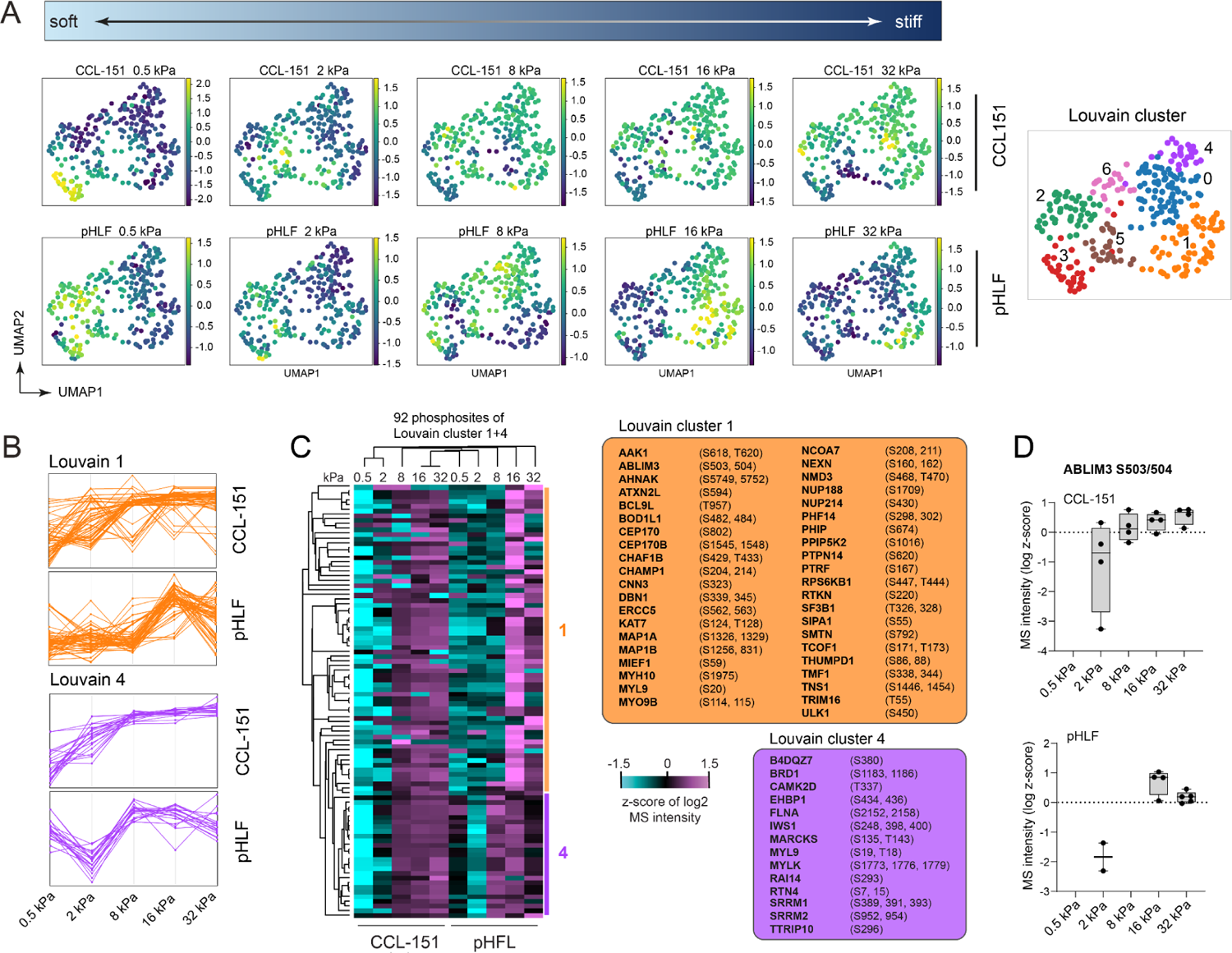
Integrative analysis of phosphoproteomes reveals conserved mechanosensitive sites in primary human lung fibroblasts. A. The UMAP shows integrated phosphosites from CCL-151 and pHLF experiments. Each dot represents one unique phosphorylation site. The color code displays phosphorylation intensities in the indicated experimental conditions. In the right panel 7 clusters of phosphosites calculated by the louvain algorithm are shown. A. The line plots illustrate the z-scored kinetic profiles of phosphosites in louvain cluster 1 and 4 for the two experiments. B. Heatmap depicts hierarchical clustering (Pearson correlation) of the z-scored MS intensities across the indicated conditions. C. The box plot shows phosphopeptide intensities of ABLIM3 S503/504 across the indicated experimental conditions.

In particular louvain cluster 1 and cluster 4 displayed comparable kinetic profiles between CCL-151 and pHLF (Figure 3B) and thus represent highly conserved mechanosensitive phosphorylation sites (Figure 3C), such as ABLIM3 S503/504 (Figure 3D), which plays a role in regulation of serum-response factor (SRF)-dependent transcription (Barrientos *et al*, 2007). We subjected sites in Louvain cluster-1 and −4 to Metascape based pathway enrichment analysis and found that cluster-1 was enriched for GO terms such as ‘actin cytoskeleton organization’ and ‘rRNA metabolic process’, as well as the reactome pathway RHO GTPases activate Citron kinase. Cluster-4 showed significant enrichment for the GO terms ‘actin cytoskeleton organization’, ‘histone modification’ and ‘regulation of cell morphogenesis’, as well as the reactome pathway ‘RHO GTPases activate PAK’. Furthermore, cluster-4 was enriched in the DisGenNET terms ‘fibromuscular dysplasia’ and ‘lung diseases’. In summary, we have identified candidate signaling pathways and kinases involved in feedback connections of lung fibroblasts sensing the stiffness of their ECM environment.

### Time-resolved analysis of lung fibroblast mechanosensing

When cells first adhere to the ECM substrate they undergo a process of cell spreading, which for CCL-151 lung fibroblasts on stiff substrates was isotropic in the first 30-60 minutes, followed by cell polarization and formation of large focal adhesions and stress fibers within the next hour (Figure 4A, B).

**Figure 4.**
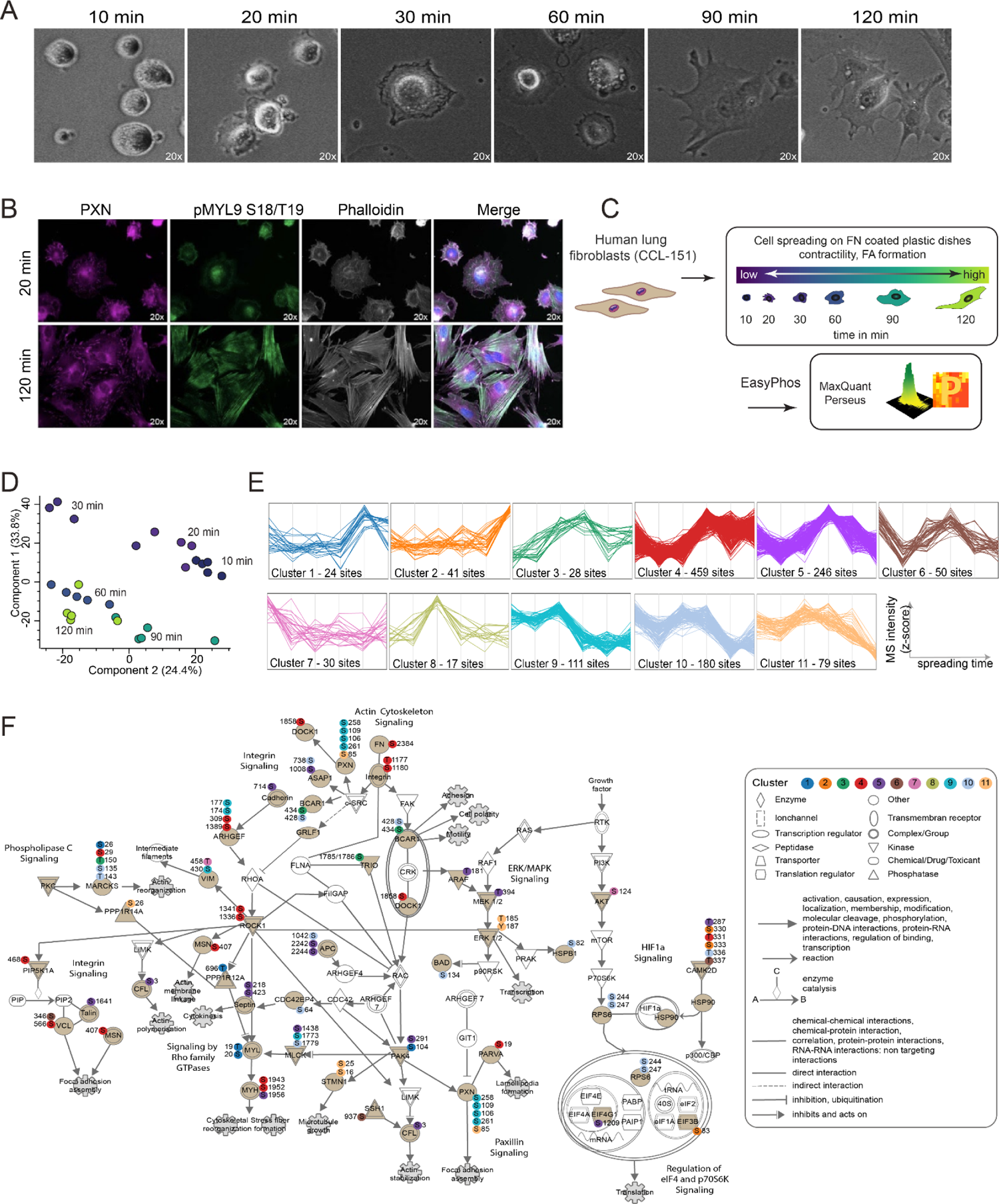
Time resolved analysis of phosphoproteome dynamics reveals distinct phases of cell spreading. A. Phase contrast images show representative CCL-151 cells seeded on fibronectin-coated glass coverslips at the indicated timepoints. B. Immunostainings show CCL-151 cells after spreading 20 or 120 minutes. Staining of PXN in magenta, phosphorylated MYL9 (Thr18/Ser19) in green and actin (Phalloidin) in white. C. Experimental design. D. PCA of the regulated phosphoproteome during cell spreading. The two first components separate the data according to spreading time. E. The 11 clusters with distinct activation profiles were defined by hierarchical clustering (Pearson correlation, FigS4F). F. Mapping of a subset of regulated phosphosites during cell spreading on canonical signaling networks. The phosphorylation sites are color-coded according to their cluster identities shown in (E). Proteins color coded in light brown are contained in the phosphoproteomic data set. The white colored proteins were either not found to be phosphorylated in the data set or measured but were excluded by the settings of the performed ANOVA test.

Increased cell contractility over time was associated with increased phosphorylation of the myosin light chain on MYL9 T19/S20 (Figure 3B). To globally reconstruct cellular signaling during cell spreading and generate a time-resolved map of mechanosensitive phosphosites we measured phosphoproteomes at 10, 20, 30, 60, 90, and 120 minutes after seeding on FN coated stiff substrates (Figure 3C). A total of 12364 class-I phosphosites were detected (Table S3). PCA analysis showed good agreement between experimental replicates and a clear separation of timepoints (Figure 4D). We detected 42.8 % singly, 44.1 % doubly and 13.1 % triply phosphorylated peptides, with serine being the most phosphorylated residue, followed by threonine and only few tyrosine phosphorylations (Figure S4A).

The data set contains 689 regulatory phosphosites with a known function, and 883 sites that were not covered in the PhosphositePlus database (Figure S4B). The average phosphoproteome depth per replicate sample was 7224 (Figure S4C). Looking into two mechanosensitive phosphosites validated in the first experiment we found gradually increased PXN S126 and MYL9 T18/S19 over time (Figure S4D, E). ANOVA analysis (1 % FDR) resulted in 1265 significantly regulated phosphosites (Figure S4F, Table S4), which comprised two main clusters, representing the early phase of cell attachment (10-30 min) and the polarized contractile phase (60-120 min) respectively (Figure 4D and S4F). This broad categorization was refined by hierarchical clustering of the sites which resulted in eleven different cluster trends (Figure 4E). To highlight potential functional connections between these sites we mapped them according to their cluster identity on distinct signaling pathways (Figure 4F), illustrating relevant sections of the pathways ERK/MAPK signaling, actin cytoskeleton signaling, integrin signaling, phospholipase C signaling, Paxillin signaling, signaling by Rho GTPases signaling, HIF1a signaling, and eiF4 and p70S6K signaling (Figure 4F). We used Metascape and Ingenuity Pathway Analysis (IPA) for prediction of regulated pathways and upstream regulators (Figure S4G-I). Out of the 1265 significant sites, we found 90 sites with known regulatory functions in cell adhesion, cytoskeletal reorganization, cell motility, cell growth and transcription (Figure S5).

To integrate the stiffness sensing and cell spreading time-course experiments, the identity of significantly regulated sites across the two datasets was compared. We identified 227 phosphorylation sites that were significantly regulated in both data sets (Figure S6A; Table S5), and visualized the overlap in a shared UMAP embedding (Figure S6B). Hierarchical clustering analysis (Pearson correlation) of this integrated dataset revealed three clusters with distinct regulation (Figure 5A). In cluster A we found 42 mechanosensitive sites that were increased on stiff substrates and peaked late in the cell attachment time-course (Figure 5B). Interestingly these sites were enriched for cytoskeletal processes on both actin and microtubule filament networks as well as the regulation of histone modifications and the ‘CDC5L protein complex’, which is essential for pre-mRNA splicing (Figure 5C).

**Figure 5.**
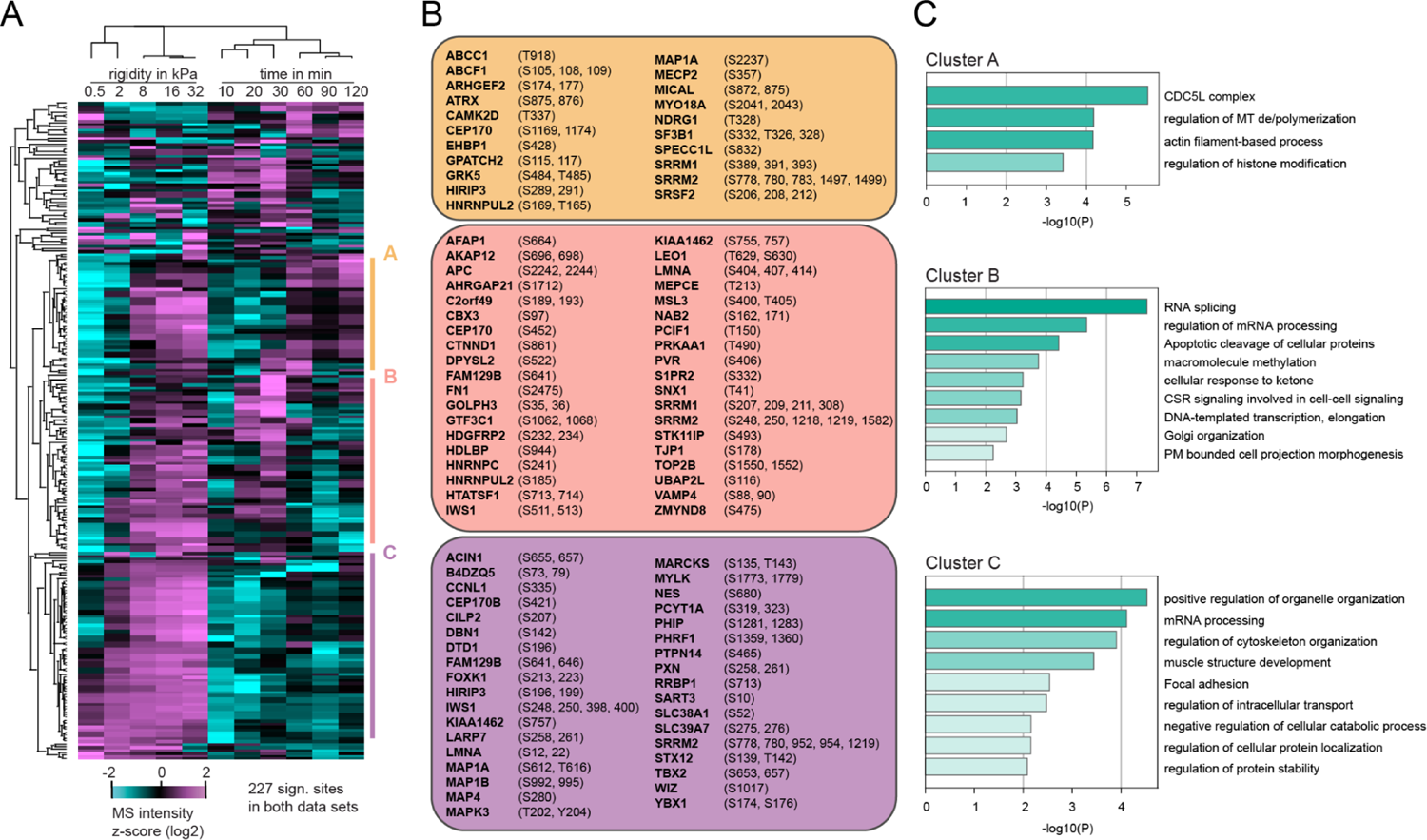
Kinetics of mechanosensitive sites after cell attachment. A. Heatmap depicts hierarchical clustering (Pearson correlation) of the z-scored MS intensities across the indicated conditions. B. Corresponding proteins to the regulated phosphosites with their respective phosphorylated sites in the clusters A, B and C. C. Bar graph shows enriched terms across gene lists of clusters A, B and C, colored by -log10 p-values. Enrichment analysis was performed with Metascape (www.metascape.org).

In cluster B we identified 61 phosphorylation sites belonging to 38 proteins, which are enriched for GO terms such as ‘RNA splicing’ and ‘regulation of mRNA processing’ as top enriched annotations (Figure 5B, C). Overall, the mechanosensitive sites in this cluster were increased on stiff substrates but in contrast to cluster A already peaked at 20–30 minutes after cell attachment. Cluster C contained 63 phosphosites corresponding to 34 different proteins. Here the increase in phosphorylation was already observed above 2 kPa and also the phosphorylation status of the corresponding sites in the spreading time course was continually increasing over time. In summary, we used time-resolved phosphoproteomics of cell spreading on a stiff fibronectin substrate to analyze the temporal dynamics of mechanosensitive signaling events.

### NFATC4 regulates collagen secretion, cell invasion and contractility

Our phosphoproteomic portrait of fibroblast mechanosensing revealed a large number of candidate kinases and transcriptional regulators that react to ECM substrate stiffness in fibroblasts. We were particularly interested in the transcriptional regulator NFATC4 (NFAT3) because, to the best of our knowledge, its regulation by mechanical forces and its relevance for tissue fibrosis has been previously unknown. We found ECM stiffness dependent phosphorylation of S213/S217 on NFATC4, which was described to induce its activity as a transcriptional regulator (Yao *et al*, 2007) (Figure 2). Therefore, we hypothesized that NFATC4 activity might be needed for the pathological activity of myofibroblasts in tissue fibrosis.

Tissue stiffness is increased during lung fibrosis, which coincides with a transition of fibroblasts into ACTA2+/CTHRC1+ contractile myofibroblasts that secrete large amounts of collagens and invade the tissue (Mayr *et al*; Tsukui *et al*, 2020). To test the functional relevance of NFATC4 in fibroblast (patho-)physiology we used siRNA to knock down *NFATC4* expression in CCL-151 (Figure 6) and pHLF from an idiopathic pulmonary fibrosis (IPF) patient (Figure S7). NFATC4 expression was consistently downregulated >70% for at least 2–3 days after siRNA transfection. Interestingly, we observed slightly reduced levels of *NFATC4* mRNA in cells plated on stiff substrates compared to soft (Figure 6A). Stiff substrates induced the expression of the myofibroblast marker alpha smooth muscle actin (encoded by the gene *ACTA2*), as well as collagen type-I (C*OL1A1*), both of which were significantly reduced in *NFATC4* knock down cells (Figure 6A), indicating that *NFATC4* indeed is involved in myofibroblast biology.

**Figure 6.**
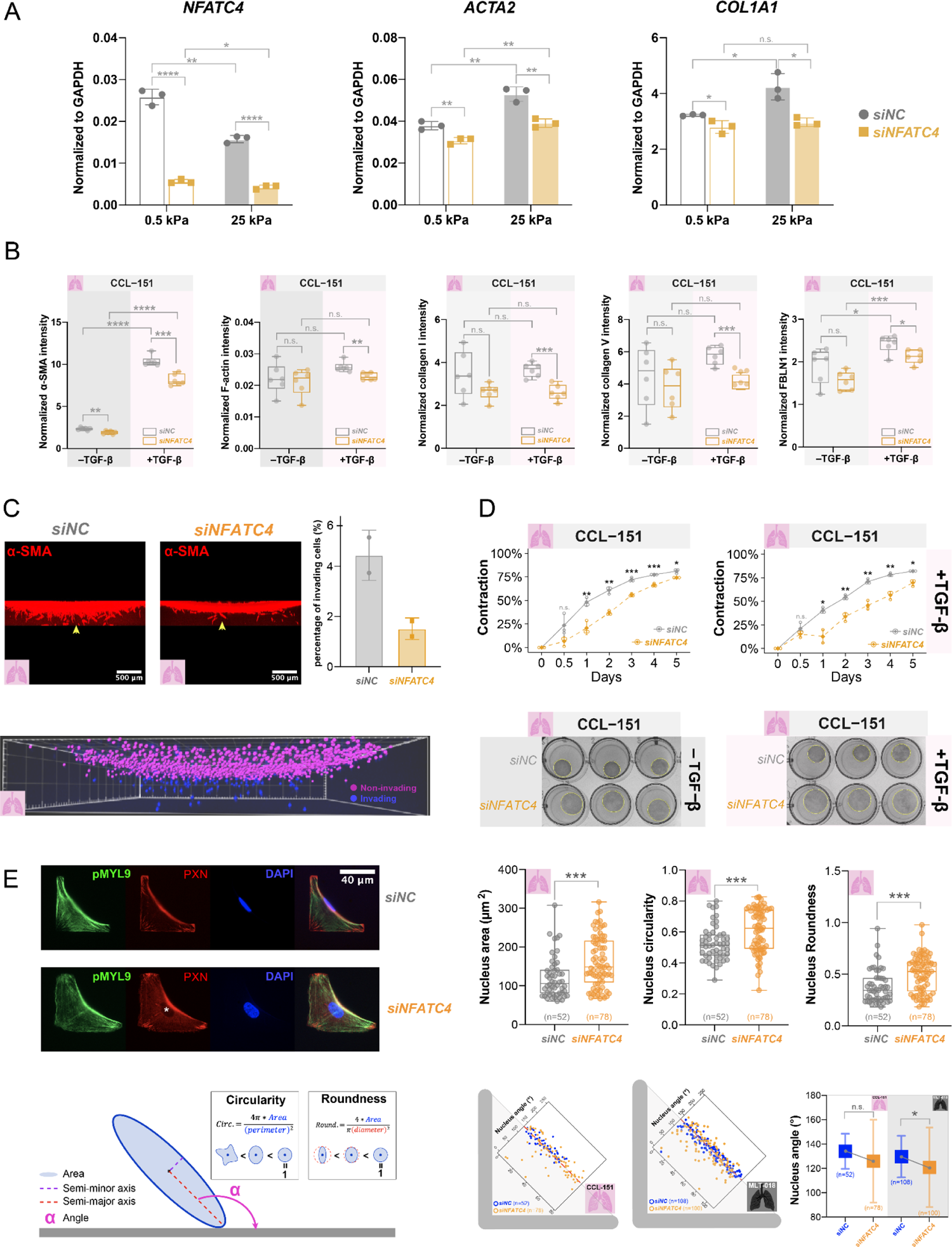
*NFATC4* knockdown attenuates fibroblast-to-myofibroblast transition, collagen deposition, invasion, and cell contractility. A. qPCR analysis of mRNA expression levels of *NFATC4*, *ACTA2* and *COL1A1* genes in CCL-151 transfected with *siNC* or *siNFATC4* on stiff and soft substrates (n = 3). B. Quantification of intracellular α-SMA, F-actin, and extracellular deposited collagen I, collagen V, and Fibulin I fluorescence intensity normalized to DAPI (n = 6). C. Collagen invasion. Representative confocal immunofluorescence images of collagen invasion assay after 3D projection (left panel), fibroblasts were stained for α-SMA (encoded by the gene *ACTA2*) (red). The yellow arrow indicates an invasive cell. Scale bar = 500 μm (top left and top middle panel). The nuclei (DAPI channel) were used to quantify the percentage of invading fibroblasts. Data were imported to the software Imaris for 3D visualization, and a representative image is shown at the bottom panel. The quantification results were visualized with bar plots, and data are expressed as relative mean ± standard deviation from two replicate experiments (right panel). D. CCL-151-induced collagen contraction assay. Contraction of collagen gels measured over different culture periods in the absence (top left panel) or presence (top right panel) of TGF-β (1 ng/mL). Each circle on the line denotes an individual contraction value (n = 3). The corresponding images of day 3. The yellow circles outline the area of collagen gel pads. E. Quantification of nuclear deformation on micropatterns. Representative fluorescent images of CCL-151 fibroblasts on micropatterned coverslips. Staining of paxillin reveals focal adhesion (red), phospho-myosin light chain 9 (p-MYL9) indicates active myosin (green), F-actin marks stress fiber (white), and DAPI stains the nuclei. Most of the nuclei of siNC fibroblasts were closer to the adhesion-free edge of the L-patterns and appeared more compact, while most of the nuclei of *siNFATC4-transfected* cells were in the center of the cells and appeared spheroid (left). The nucleus area, circularity, and roundness were quantified (top right). The related formula is shown at the bottom left. The nucleus angles were also measured and presented as scatter plots, with the x-axis representing the cell index (ranging from 1 to the total number of cells in the group), and the y-axis representing the nucleus angle (dashed red line indicates a value of 135). The average angles were compared (bottom), Icons: pink lung: CCL-151; black lung: MLT-018. Abbreviations used in the formulae: Circ., Circularity; Round., Roundness. The area and roundness of fibroblast nuclei were quantified based on DAPI staining (right) using ImageJ. Statistics: *p < 0.05, **p < 0.01, ***p < 0.001, ****p < 0.0001, and n.s.: not significant according to the t-test.

In addition to qPCR based testing of gene expression we analyzed *ACTA2* and F-actin levels as well as the expression, secretion and assembly of type-I (C*OL1A1*), type-V collagen (*COL5A1*), Fibulin-I (FBLN1), all of which are key hallmarks of myofibroblast identity, using antibody based immunofluorescence analysis (Figure 6B). Cells were plated on stiff fibronectin coated tissue culture plates and treated with transforming growth factor beta (TGF-β) to boost fibroblast to myofibroblast conversion. Indeed, we observed a significant upregulation of *ACTA2* upon treatment with TGF-β (Figure 6B). The *NFATC4* knock down significantly reduced the amount of F-actin as well as protein amounts of *ACTA2, COL1A1, COL5A1,* and *FBLN1* under these conditions in both CCL-151 (Figure 6B) and IPF derived pHLF (Figure S7, S8). ACTA2+/CTHRC1+ myofibroblasts have been shown to be highly invasive *in vivo* and *in vitro* (Mayr *et al*; Tsukui *et al*, 2020). We seeded CCL-151 fibroblasts on top of collagen gels and assessed the proportion of invading cells using confocal imaging in z-stacks. In the course of 96 hours, significantly fewer NFATC4 knock down cells invaded the 3D collagen gel (Figure 6C).

Next, we asked if the ability of fibroblasts to contract a collagen type-I spheroid was affected by *NFATC4* expression levels. We seeded 5×10^4^ CCL-151 fibroblasts (Figure 6D) or MLT-018, a IPF-patient derived pHLF (Figure S7) into collagen gels and analyzed the contraction of the gels over time. *NFATC4* knock down cells showed significantly reduced levels of gel contraction in both CCL-151 and IPF derived pHLF (Figure 6D).

To analyze the effect of NFATC silencing on the cell contractility feedback on stiff substrates in single cells we plated siRNA transfected cells on fibronectin coated adhesive micropatterns with an L shape. In this system, the cells adhere to the L-shape and form a contractile actomyosin bundle spanning the non-adhesive edge of the L-pattern. Cytoskeleton-mediated contraction has been shown to be required for the fibroblast-to-myofibroblast transition (Meyer-ter-Vehn *et al*, 2006) and can also affect nuclear morphology (Alisafaei *et al*, 2019). Interestingly, it has been shown that fibroblasts activated by increased stiffness have less round nuclei (Walker *et al*, 2021). We therefore analyzed the nuclear shapes and distributions in single fibroblasts.

We noticed that most nuclear centroids of *siNC*-transfected fibroblasts were pulled towards the contractile adhesion free edge on the L-shaped micropatterns (Figure 6E, Figure S10A, Figure S10C), which also resulted in an elongated spherical nuclear shape (Figure 6E, Figure S10B). In contrast, the nuclei of *siNFATC4*-transfected cells were distributed more widely, with a tendency to be positioned away from the contractile edge and oriented more towards the adhesive edge (Figure 6E, Figure S10A). The nuclei of *siNFATC4*-transfected cells displayed a significantly larger area and had a relaxed round shape (Figure 6; Figure S10), suggesting an overall reduction of cytoskeletal and nuclear tension in *NFATC4* knock down cells. F-actin stress fibers decorated with active myosin (pMLC S19/S20) were predominantly located at the contractile non-adhesive edge in *siNC*-transfected, while these fibers were more widely distributed in *siNFATC4*-transfected fibroblasts (Figure 6E, Figure S10A).

In conclusion, our loss of function data reveals a novel function of the transcriptional regulator NFATC4 in myofibroblast biology, including fibroblast-to-myofibroblast transition, cell contraction, ECM deposition, as well as collagen invasion.

## Discussion

The growing field of mechanobiology, at the interface of physics, biology and bioengineering, established that forces applied to cells as well as physico-chemical properties of the cellular microenvironment affect almost all aspects of multicellular life. Cellular responses to mechanical forces are substantial in embryonic development and adult physiology and are involved in many diseases, such as atherosclerosis, hypertension, osteoporosis, muscular dystrophy, myopathies, cancer, metastasis formation and fibrosis (Hoffman *et al*, 2011; Dupont & Wickström, 2022). The molecular details of these mechanosensitive signaling cascades are however still not very well characterized on a proteome wide scale. In this work we used label free mass spectrometry driven phosphoproteomics to quantify thousands of signaling intermediates during the process of cellular mechanosensing. Our data (1) reveals a major cell intrinsic stiffness threshold for mechanosignaling in primary human lung fibroblasts (2–8 kPa), above which the cells undergo a fibroblast to myofibroblast transition seen in fibrotic disease. We (2) predict kinases, signaling pathways and transcriptional regulators that are associated with this mechanosensitive cell state transition. We use A. (3) time-resolved analysis to study the kinetics of mechanosensitive phosphorylation events, and (4) establish the functional relevance of the transcriptional regulator NFATC4 during this process.

Which cellular structures mediate mechanosensing? Recent data indicates that multiple organelles, including the plasma membrane, nucleus, mitochondria and ER, display mechanosensitivity (Dupont & Wickström, 2022). In this work, we observed stiffness dependent phosphorylation of proteins in all of these organelles as a resource for the research field. Furthermore, contractile cytoskeletal units, cytoskeleton-binding proteins, and integrin- and cadherin-based adhesion complexes, are of key importance as structural interface to the environment. We and others pioneered the use of mass spectrometry driven proteomics to systematically analyze the molecular composition of the cell-matrix adhesome and its dynamic changes under the influence of mechanical tension (Geiger & Zaidel-Bar, 2012; Horton *et al*, 2015; Humphries *et al*, 2009; Kuo *et al*, 2011; Schiller *et al*, 2011; Schiller & Fässler, 2013; Schiller *et al*, 2013). Our previous results on fibroblast mechanosensing revealed that β1-family integrins and associated focal adhesion proteins feed into signaling pathways that produce myosin-II mediated force when bound to fibronectin, while the αV-family of integrins and specific associated focal adhesion proteins respond to myosin-II and matrix stiffness dependent tension at focal adhesions to reinforce the adhesion site. Both systems act in synergy to enable mechanoreciprocal reinforcement of cell contractility while sensing ECM stiffness (Schiller et al, 2011; Schiller et al, 2013a). Our new data features a large number of cell-matrix adhesome associated and cytoskeletal proteins that are differentially phosphorylated during mechanosensing, which forms the basis for future functional studies of individual phosphoproteins or phosphorylation sites during mechanosensing.

We also uncover the mechanosensitive regulation of a large number of phosphorylation sites on transcriptional regulators, including FOXO3, MAF1, YBX1, and NFATC4, thereby potentially linking the immediate feedback connections in the cytoskeletal apparatus with decisions on long term gene expression and cell state transitions. NFATC4 (NFAT3) is a Ca^2+^-regulated transcription factor that is involved in several processes, including the development and function of the immune, cardiovascular, musculoskeletal, and nervous systems (Hoey *et al*, 1995; Yang *et al*, 2001, 2002; Yao *et al*, 2007; Cho *et al*, 2007; Qin *et al*, 2008; Mei *et al*, 2015). We find the NFATC4 S213/S217 site induced by substrate stiffness, suggesting increased transcription factor activity on stiff substrates as previously shown for these sites (Yao *et al*, 2007).

Calcium mediated signaling has been previously shown to be involved in mechanosensing in various cell types and biological systems. For instance, calcium signaling has been shown to mediate the biphasic mechanoadaptive response of endothelial cells to cyclic mechanical stretch (Miroshnikova *et al*, 2021). Cell spreading on stiff substrates induces endoplasmic and nuclear calcium release (Itano *et al*, 2003), and the ability of cellular nuclei to sense mechanical forces is at least partially dependent on calcium store release as a downstream mediator (Enyedi *et al*, 2016). Nuclear deformations reflect the balance between the mechanical properties of the nucleus and the mechanical forces mainly transmitted by cytoskeletal networks (Kalukula *et al*, 2022). We observed that the nuclei of NFATC4-knockdown fibroblasts became more relaxed on micropatterns (Figure 6E and S10), indicating that NFATC4 may regulate the expression of cytoskeleton-related genes, for example, myosin heavy chain genes, and thereby affect cytoskeletal tension and mechanotransduction from ECM to nucleus.

The role of the Ca^2+^-NFATC4 axis in lung fibroblast mechanosensing is novel, but the calcium-calcineurin-NFAT signaling has been indicated in the regulation of the cytoskeleton in different cell types. For example, in myoblasts the expression of different myosin heavy chain genes depended on different NFAT isoforms (Allen *et al*, 2001; Calabria *et al*, 2009; Meissner *et al*, 2011). A recent analysis of liver fibrogenesis provides interesting data that supports our conclusions. In this work the expression of RCAN1, which inhibits calcineurin-dependent transcriptional responses by binding to the catalytic domain of calcineurin A, is shown to repress the nuclear translocation and activity of NFATC4, which interestingly alleviated liver fibrosis in the CCl4 model (Pan *et al*, 2019). Together with the functional evidence provided here in primary human lung fibroblasts we propose that further analysis of the Ca^2+^-NFATC4 axis may be of therapeutic relevance in fibrotic diseases.

We acknowledge the limitations of our work. The heterogeneity of mechanosignaling across different experiments and cell lines (CCL-151 vs pHLF) was quite high, suggesting a non trivial amount of stochasticity in response as well as potential differences in cell intrinsic thresholds of mechanosignaling for different cell lines. The range of 8-16 kPa was however highly consistent as a major threshold for lung fibroblasts to activate their stiffness response, with several events already at two kPa in both CCL-151 and pHLF. It will be thus interesting to compare the cell intrinsic mechanoresponse signaling thresholds from fibroblasts inhabiting tissues with different baseline stiffness to potentially learn how such mechanosignaling thresholds are programmed and maintained.

In summary, we provide a proteome wide analysis of phosphorylation events during human lung fibroblast mechanosensing, which reveals novel kinases, signaling pathways and transcriptional regulators involved in sensing the stiffness of the cellular microenvironment. To illustrate the value of this resource we have validated the functional relevance of the transcriptional regulator NFATC4 for stiffness induced myofibroblast activity.

## Supplementary figures

**Figure S1.**
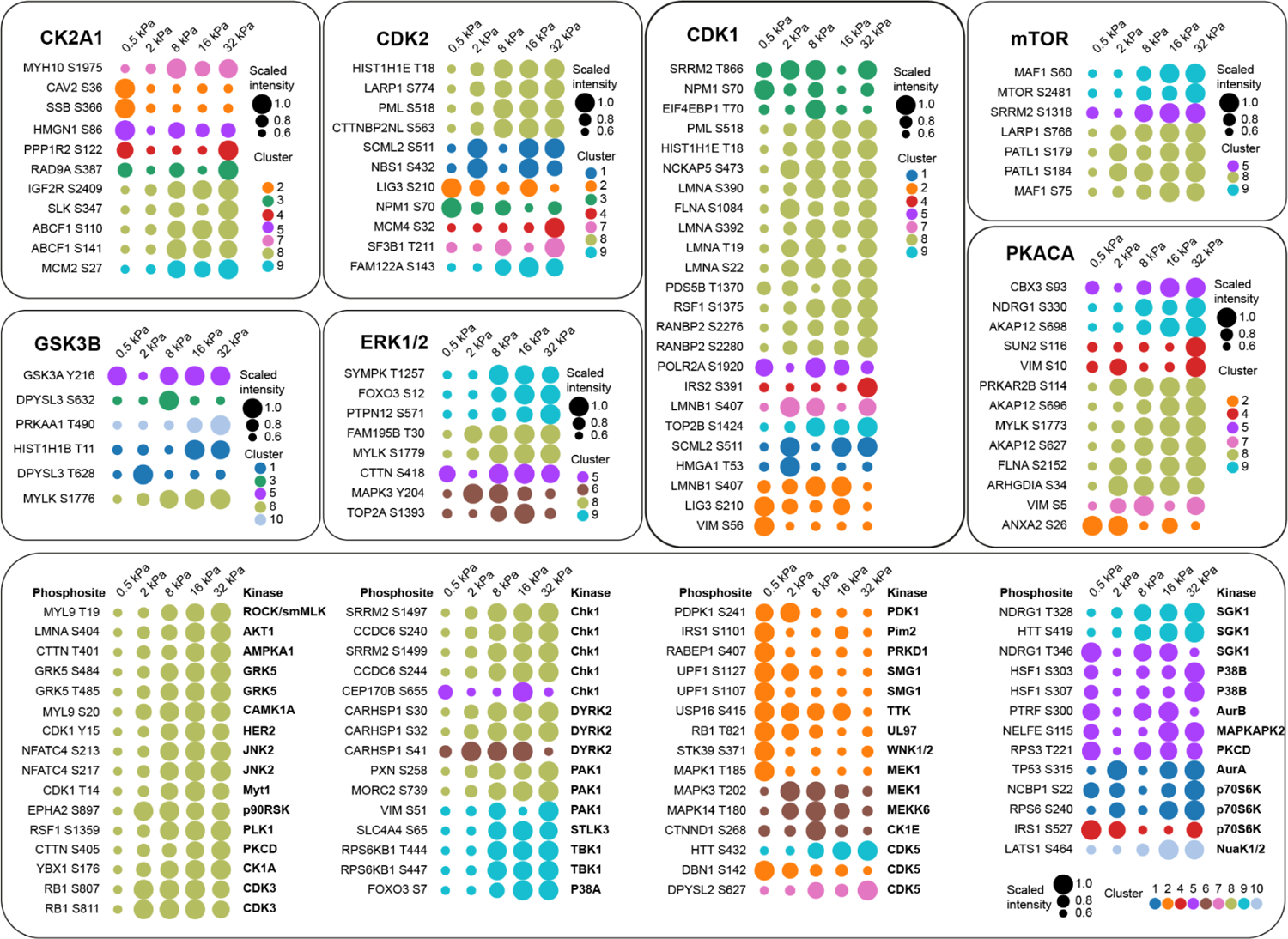
Predicted upstream kinases of mechanosensitive phosphosites. Dot plots show the dynamics of scaled phosphorylation intensities on soft to stiff substrates for selected phosphosites with known regulatory site processes (PhosphositePlus database) associated with the indicated functional categories. Cluster identities from main Figure 2 are indicated by colors. The predicted upstream kinases for the regulated motifs are shown.

**Figure S2.**
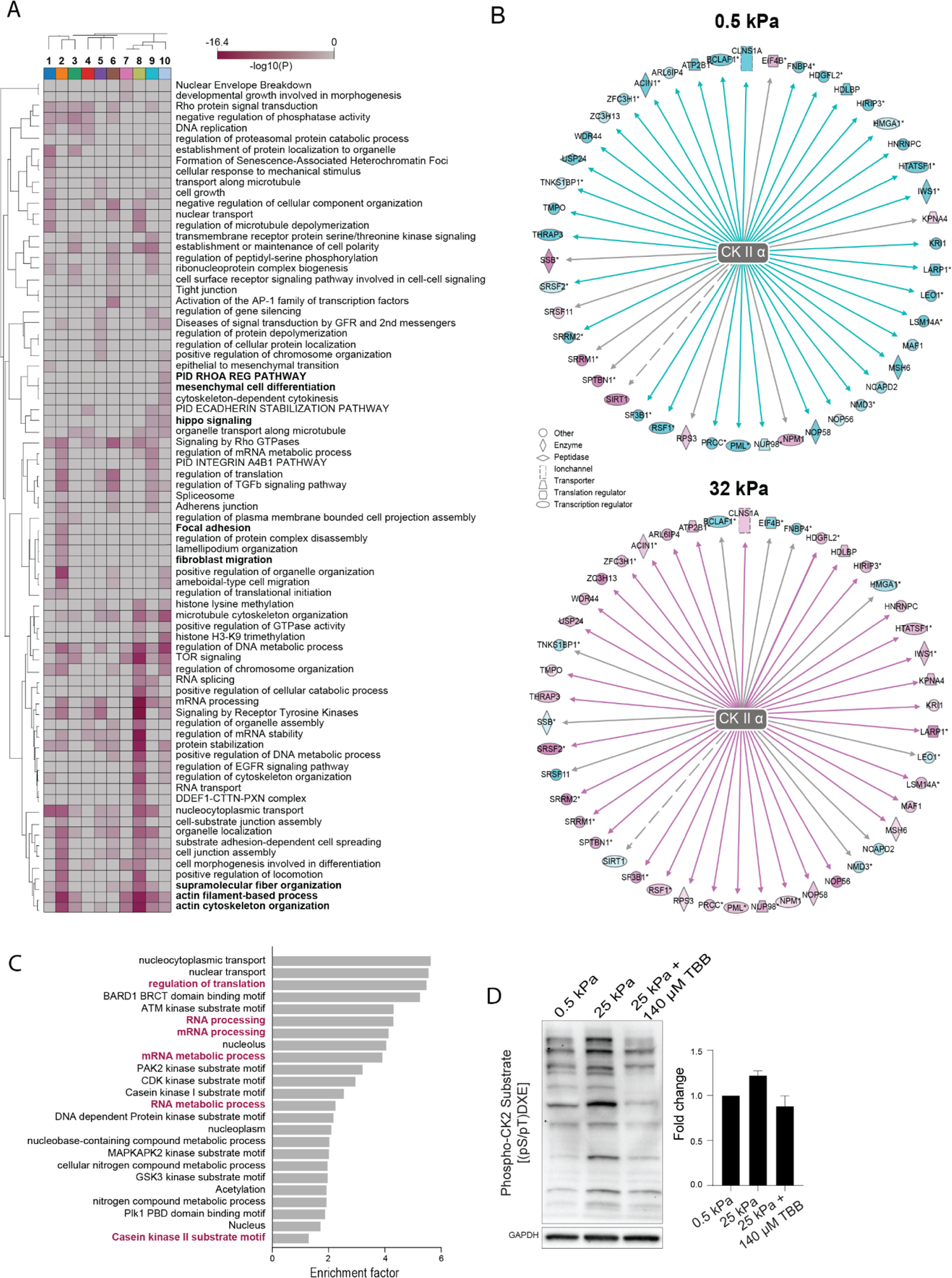
Pathway and upstream regulator analysis reveals stiffness regulated processes. A. Heat map shows enriched terms across gene lists of clusters 1–10 in main Figure 2, colored by p-values. Enrichment analysis was performed with Metascape (www.metascape.org). B. The graphs show CK2A1 targets predicted by Ingenuity Pathway Analysis (IPA) and their phosphorylation status in the phosphoproteomic data set. Turquoise color shades indicate low phosphorylation and pink color shades indicate high phosphorylation. Doted lines indicate an indirect interaction and continuous lines specify a direct interaction from prior knowledge databases. C. Results of pathway enrichment analysis of the IPA CK2A1 target sites using Fisher exact test (FDR < 5%). D. Immunoblot probed with an antibody specific for proteins containing a pS/pTDXE motif, which represents a CK2 phosphorylation consensus sequence. GAPDH served as a loading control (n=3).

**Figure S3.**
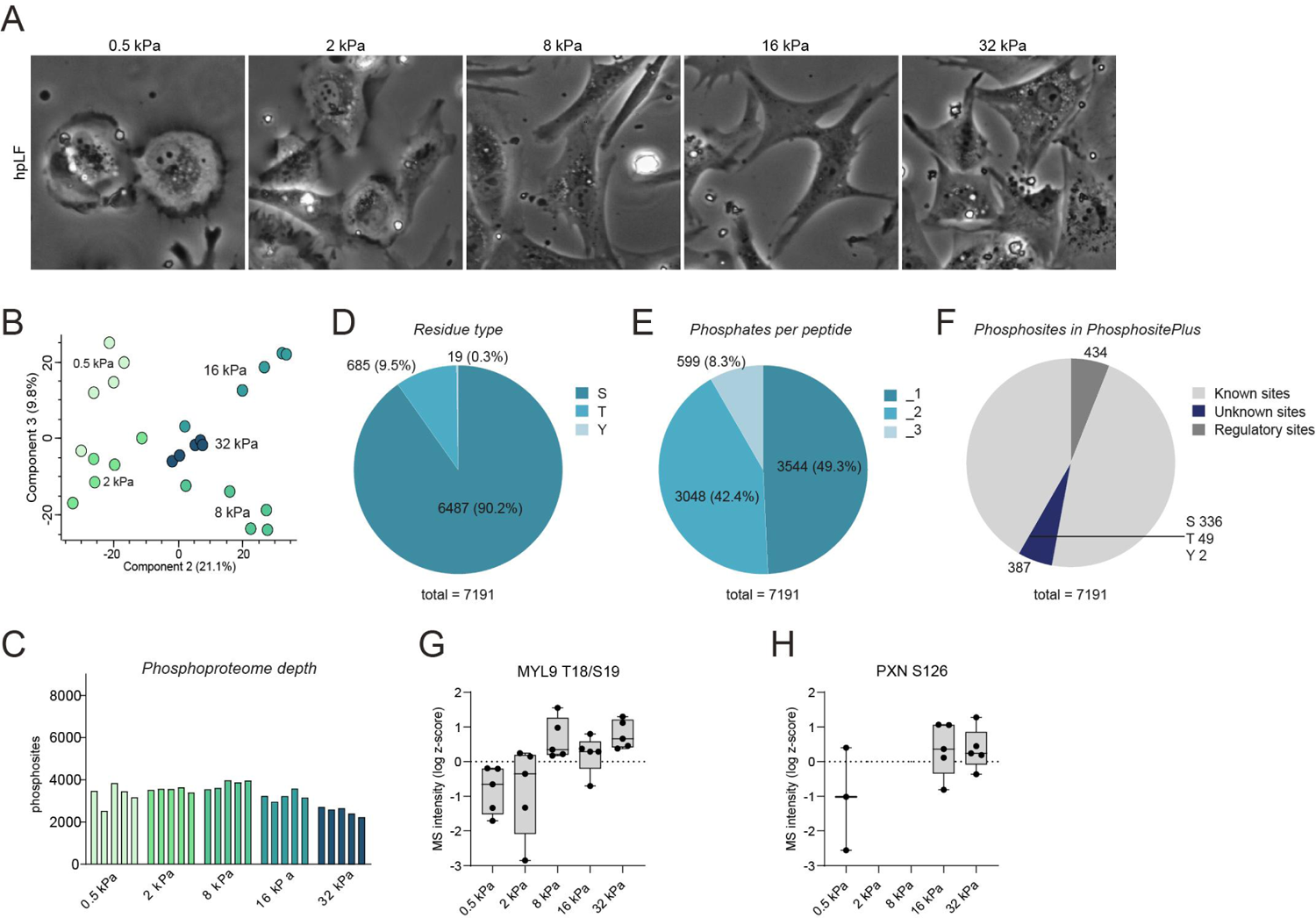
Human primary lung fibroblasts activate mechanosensitive signaling pathways. A. Phase contrast images show primary human lung fibroblasts (pHLF) from peritumoral resection of a non-CLD patient seeded on fibronectin coated substrates with the indicated stiffness. B. Principal component analysis (PCA) of the stiffness regulated phosphoproteome in pHLF. C. The bar graph shows the number of quantified phosphosites in the indicated experimental conditions. D. Pie charts represent the (D) percentage of identified phosphorylated serines (S), threonines (T) and tyrosines (Y) residues. E. The percentage of identified singly, doubly and triply (or more) phosphorylated peptides. F. Fhe number of already known sites, previously unreported sites and known sites with an assigned regulatory function in the PhosphoSitePlus database (www.phosphosite.org). G. The box plot shows phosphopeptide intensities of MYL9 T18/S19 at the indicated experimental conditions in pHLF. H. The box plot shows phosphopeptide intensities of PXN S126 at the indicated experimental conditions in pHLF.

**Figure S4.**
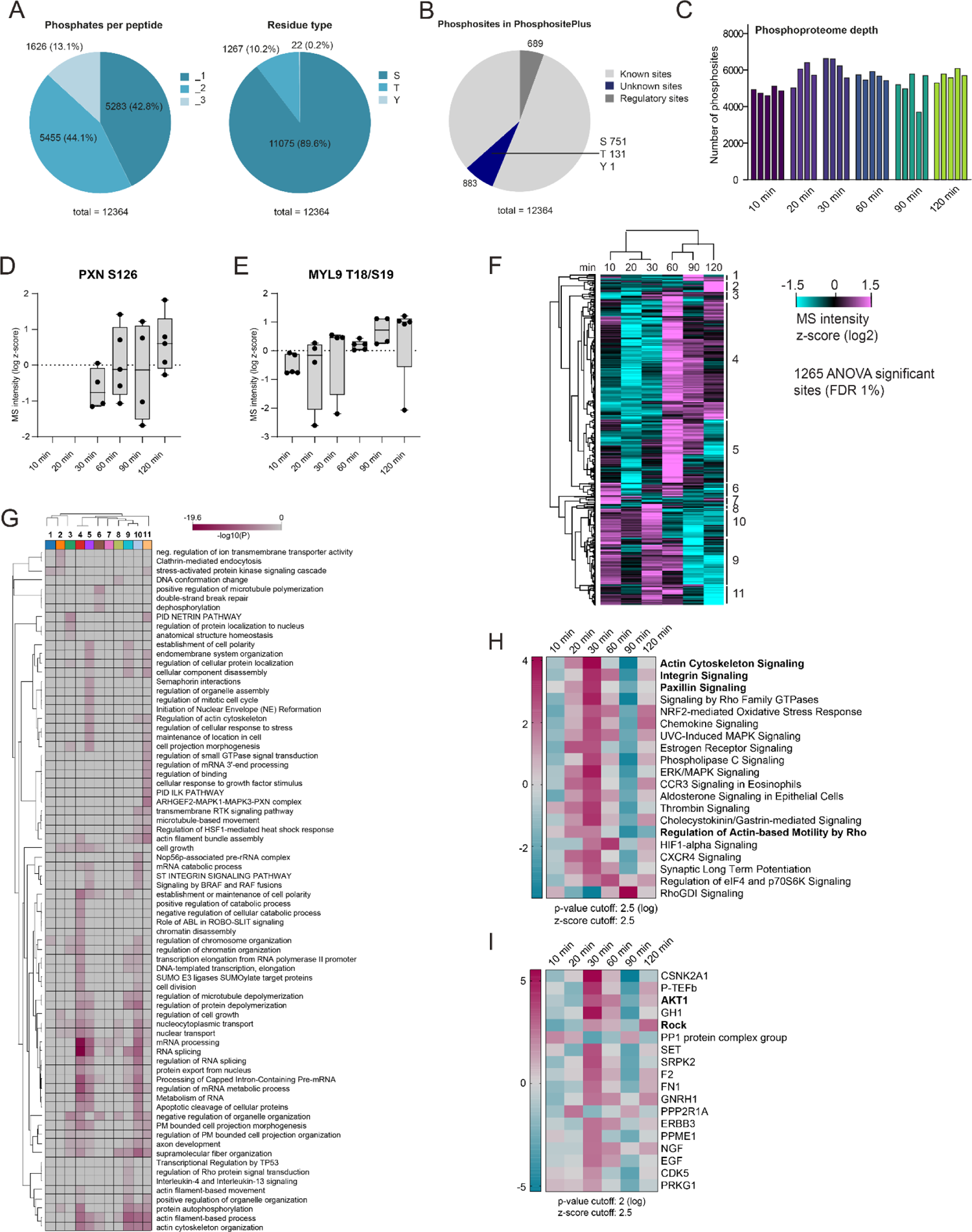
Distinct phases of cell signaling during fibroblast cell spreading on stiff substrates. A. Right panel: pie chart presenting the percentage of identified phosphorylated serines (S), threonines (T) and tyrosines (Y) residues. Left panel: pie chart presenting the percentage of identified singly, doubly and triply (or more) phosphorylated peptides. B. Pie chart showing the number of already known sites, previously unreported sites and known sites with an assigned regulatory function in the PhosphoSitePlus database. C. The bar graph shows the number of quantified phosphosites in the indicated experimental conditions. D. Phosphorylation of PXN on serine residue 126 is increasing during cell spreading on stiff substrates. E. Phosphorylation of MYL9 on threonine residue 18 and serine residue 19 is increasing with spreading time on stiff substrates. F. Heatmap shows hierarchical clustering (pearson correlation) of z-scored phosphosite intensities across the indicated timepoints. G. Heat map shows hierarchical clustering of -log10 p-values of enriched terms for the clusters in (F). Enrichment analysis was performed with Metascape (www.metascape.org). H–I. Phosphoproteomic data was analyzed using the Ingenuity pathway Analysis (IPA) platform. Significance of term enrichment is indicated through shading as log p-value. Regulated pathways belonging to “Canonical pathways” (H) and predicted “Upstream regulators” (I).

**Figure S5.**
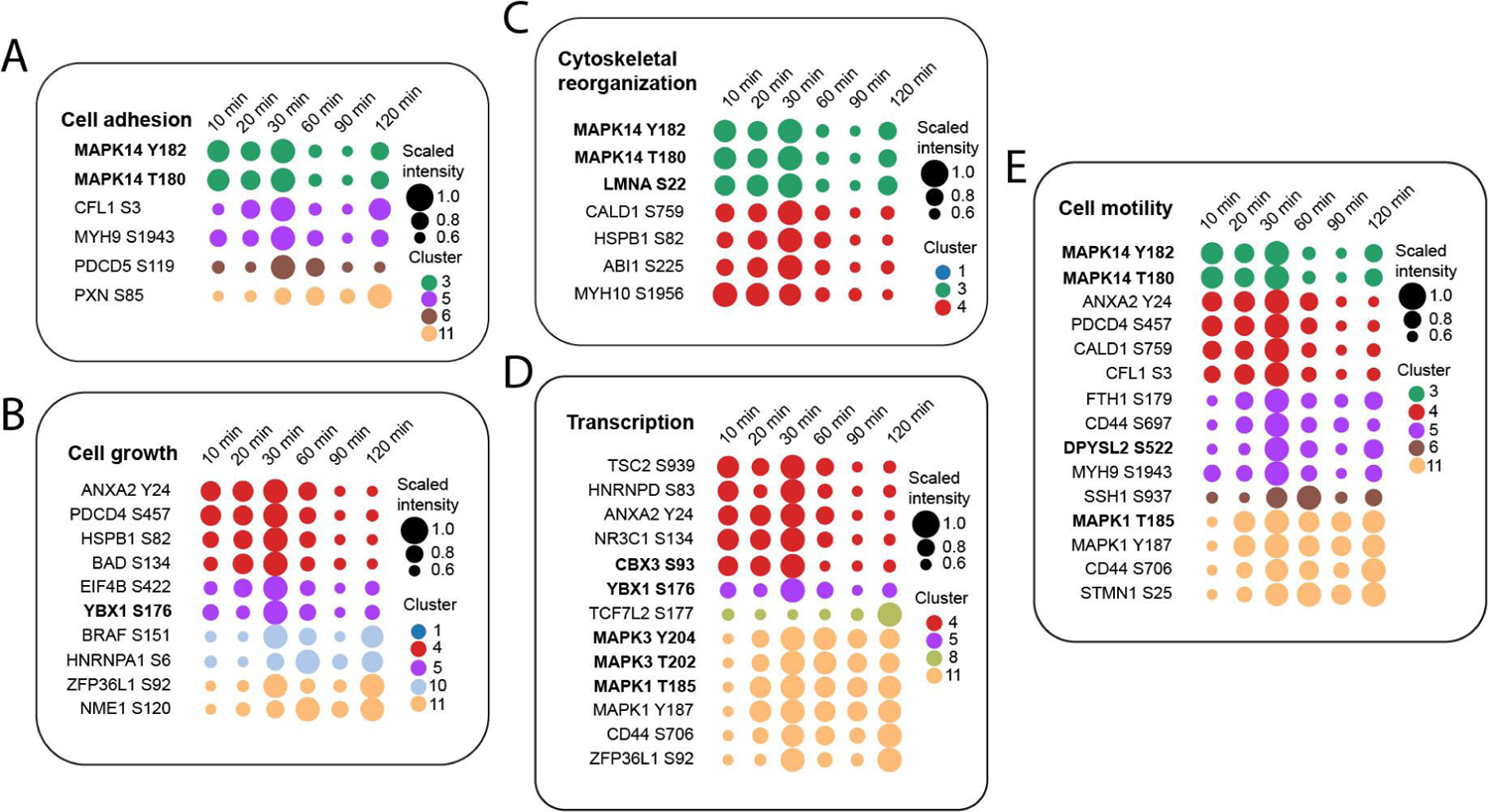
Course of phosphorylation of known regulatory phosphosites during cell spreading. A–E. Dot plots showing the dynamics of scaled phosphorylation intensities over spreading time of selected phosphosites associated with the regulatory sites process (A) ‘Cell adhesion’, (B) ‘Cell growth’, (C) ‘Cytoskeletal reorganization’, (D) ‘Transcription’ and (E) ‘Cell motility. Mechanosensitive sites are highlighted in bold font. Dots are color-codes per cluster as presented in Figure 4.

**Figure S6.**
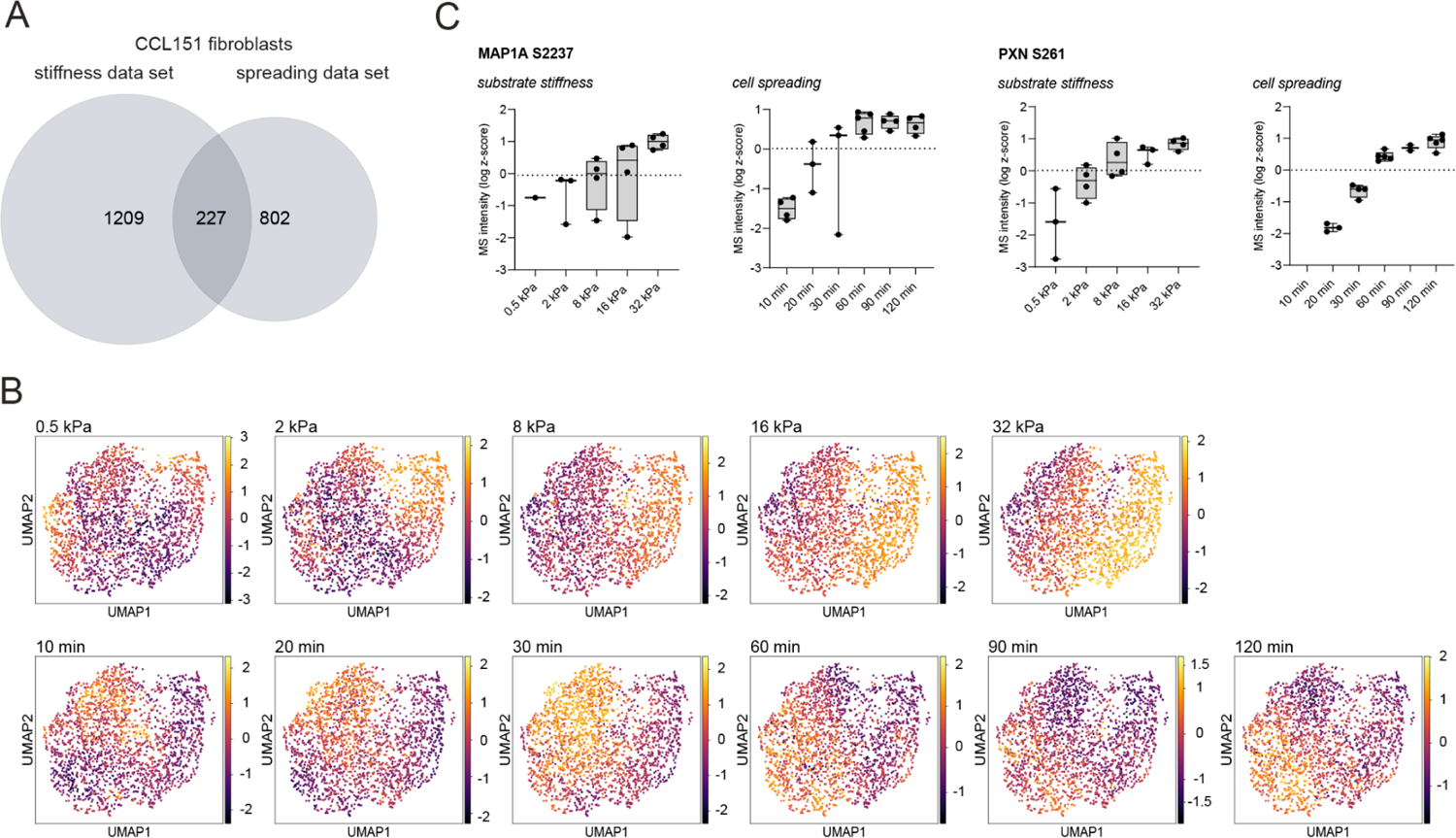
Data integration to assess time-resolved aspects of stiffness sensing. A. The venn diagram depicts the overlap of significantly regulated phosphorylation sites in both datasets. B. The UMAPs for each substrate rigidity and spreading timepoint are color-coded according to the phosphorylation intensity of each site and visualize the course of phosphorylation across both datasets. C. Box plot of phosphopeptide intensities of MAP1A S2237 and PXN S126 at the indicated experimental conditions.

**Figure S7.**
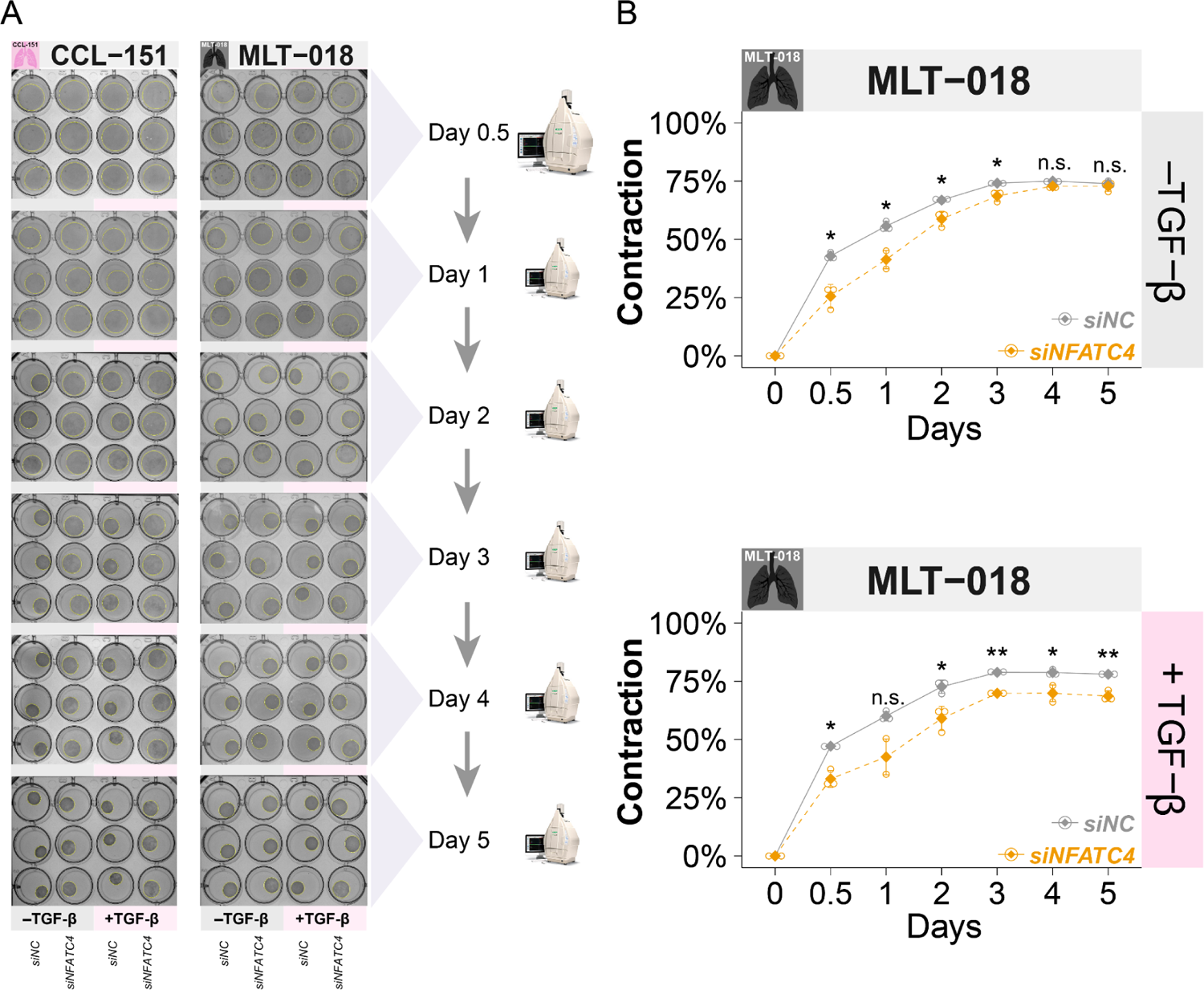
CCL-151 and MLT-018 induced-collagen gel contraction A. Pictures of CCL-151 (left) and MLT-018 (right) induced-collagen I gel contraction on days 0.5, 1, 2, 3, 4, and 5, with the absence or presence of 1ng/mL TGF-β. The area of collagen pads was outlined by the yellow circle. Areas were accordingly measured with ImageJ. B. MLT-018-induced collagen I contraction kinetics over time, with (upper panel) or without (lower panel) 1ng/mL TGF-β. Statistics: *p < 0.05, **p < 0.01, ***p < 0.001, ****p < 0.0001, and n.s.: not significant according to the t-test.

**Figure S8.**
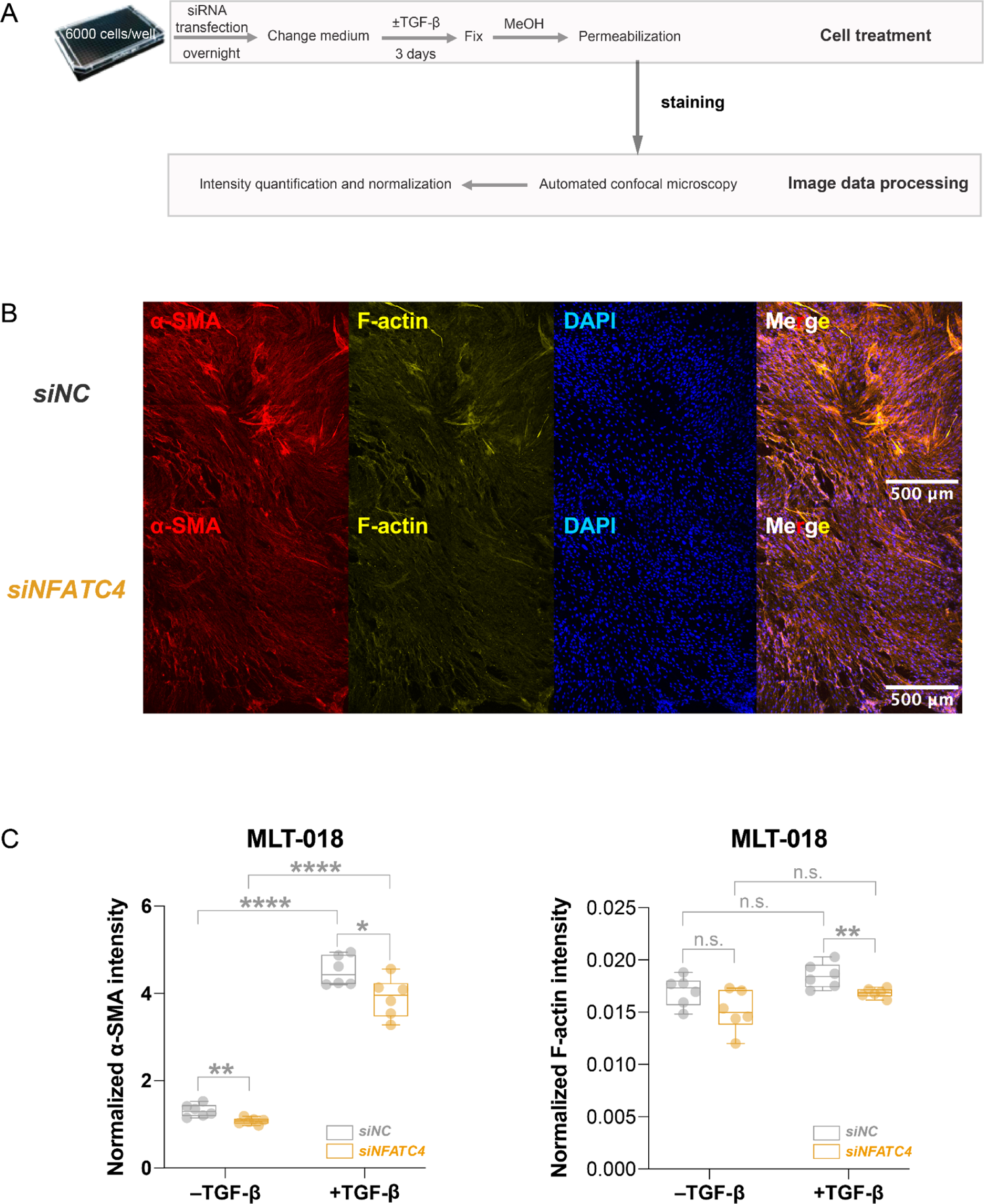
NFATC4 knockdown reduced α-SMA and stress fiber. A. Flowchart of actin staining. Fibroblasts were seeded in a Fibronectin-coated 384-well CellCarrier plate, transfected with siRNAs, treated with TGF-β (1 ng/mL in medium containing 1% FBS) for 3 days after starvation. Fibroblasts were then fixed with MeOH and stained for actin. Confocal images were acquired, and maximum intensity projection (MIP) images were generated and processed. B. Representative images showing the expression of alpha-smooth muscle actin (α-SMA) and filamentous actin (F-actin) in MLT-018 cells without TGF-β treatment. Scale bars, 500 μm. C. Quantification of expression levels of α-SMA and F-actin (fluorescence intensity normalized to Hoechst 33342 intensity) in MLT-018 cells, student’s t-test (n=6). Statistics: *p < 0.05, **p < 0.01, ***p < 0.001, ****p < 0.0001, and n.s.: not significant according to the t-test.

**Figure S9.**
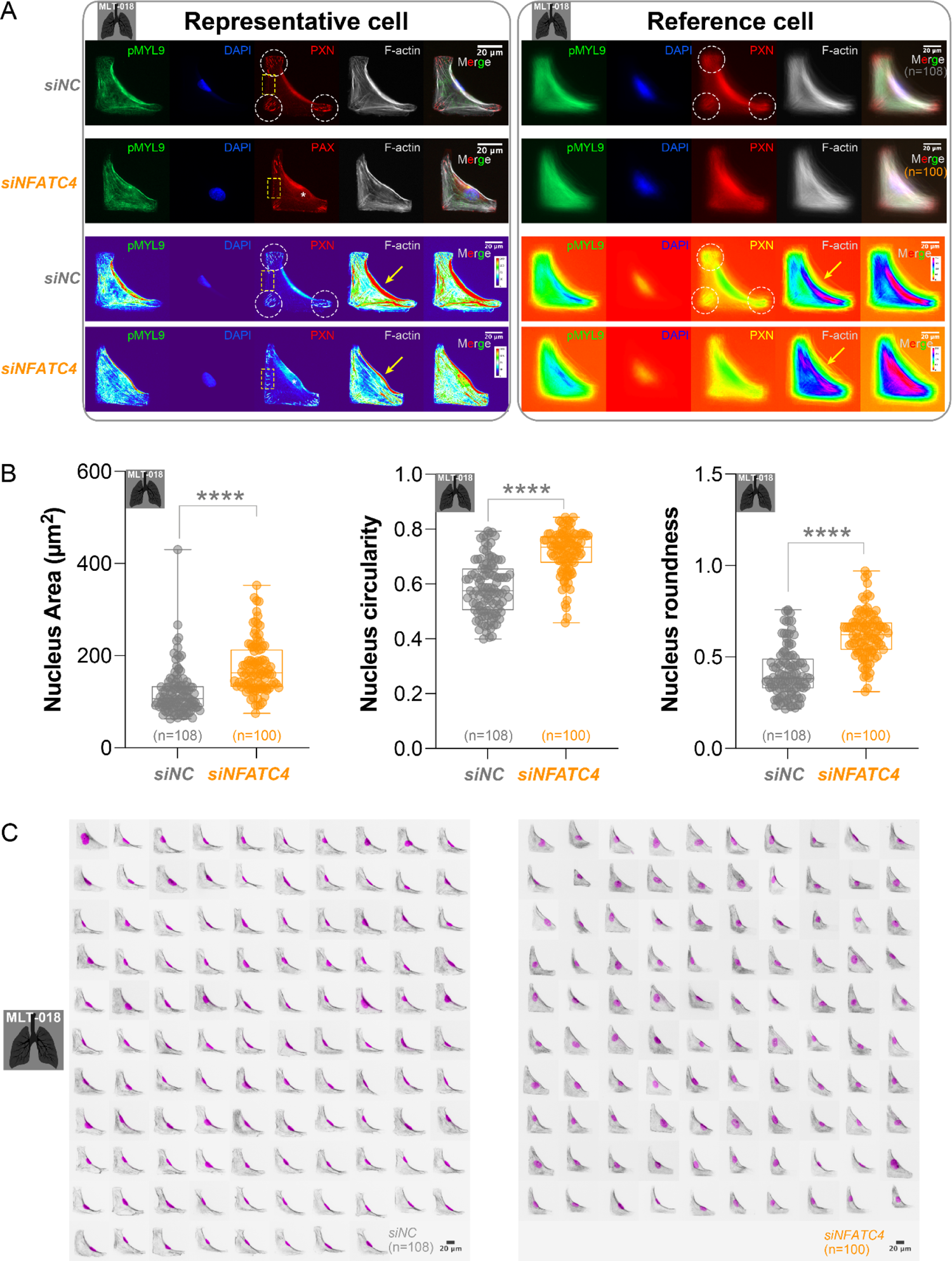
*NFATC4* knockdown reduced ECM deposition A. Flowchart of extracellular matrix (ECM) deposition assay. Fibroblasts were seeded in an Fibronectin -coated 384-well CellCarrier plate, transfected with siRNAs, and treated with TGF-β (1 ng/mL in medium containing 1% FBS) for 3 days after 24h starvation. Fibroblasts were then stained for ECM proteins. Confocal images were acquired, and maximum intensity projection (MIP) images were generated and processed. B. Representative images showing the expression of collagens I,V and FBLN1 in CCL-151. Scale bars, 500 μm. Abbreviation: Rep, Replicate. C. Quantification of expression levels of collagens and FBLN1 (fluorescence intensity normalized to Hoechst 33342 intensity), student’s t-test (n=6).

**Figure S10.**
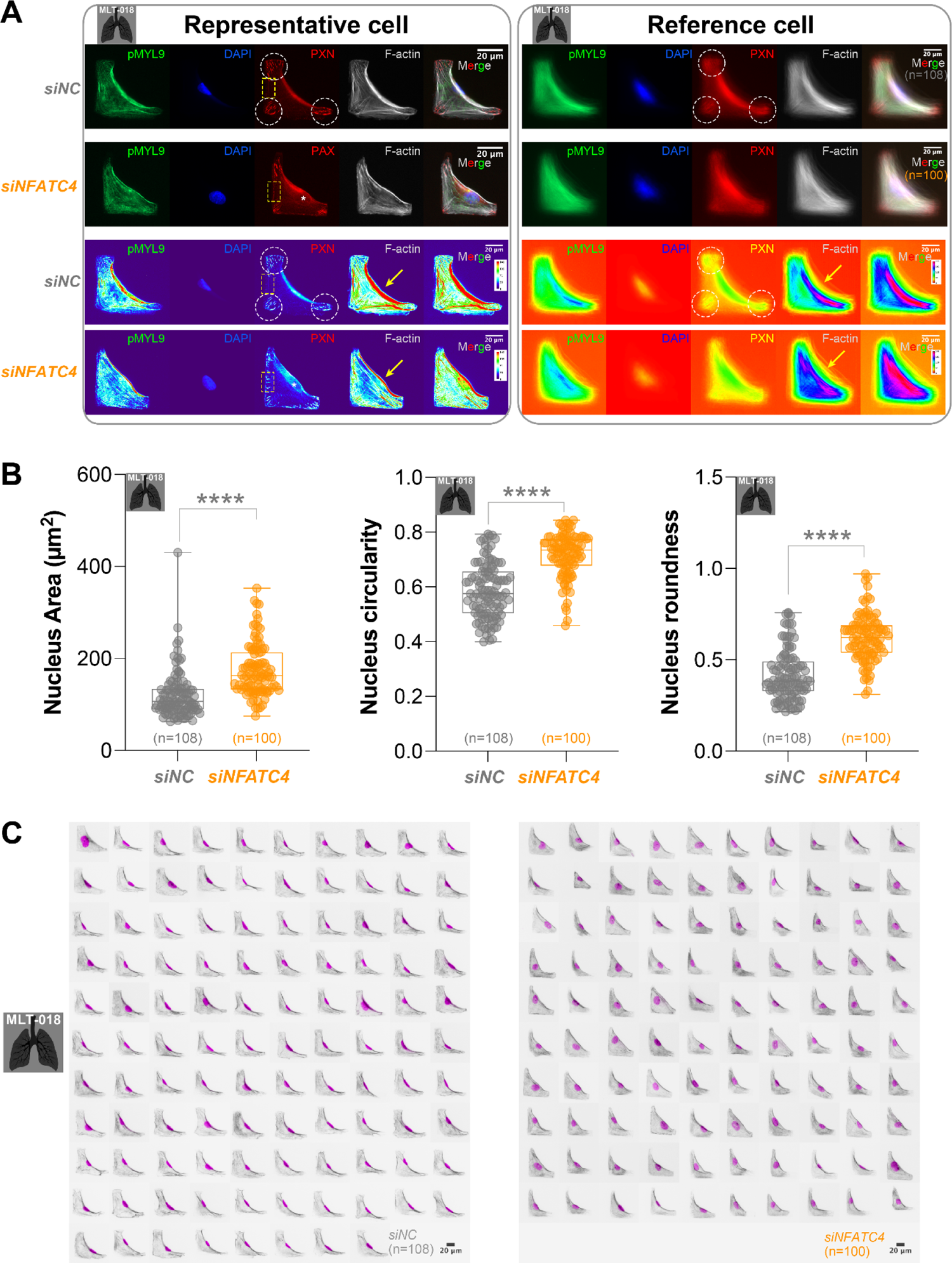
siNFATC4 affected cytoskeleton and nucleus phenotype. A. MLT-018 cells on L-shaped micropattern. were for pMYL9 (myosin activity marker, green), Paxillin (a component of focal adhesions, red), F-actin (stress fiber, gray), and nucleus (blue). A total of 108 *siNC*-transfected MLT-018 fibroblasts and 100 *siNFATC4*-transfected fibroblasts were analyzed. The Reference cell pictures (right panel) were generated using the average intensity (upper two rows). The fluorescent intensity of each channel was also illustrated by heatmaps (lower two rows). In *siNC*-transfected fibroblast, the focal adhesions indicated by Paxillin preferentially enriched at the 3 points of an L-shape micropattern (white dashed circles), while in *siNFATC4*-transfected cells, the focal adhesions also distribute on the adhesive edges of the micropattern (yellow dashed box outline, left panel). In addition, a region of the lower expression level of Paxillin was shown (white asterisk); interestingly, we found a “paxillin outline” around the nucleus in most of the *siNFATC4*-transfected MLT-018, suggesting Paxillin is associated with nucleus shape, and NFATC4 may also regulate the perinucleus Paxillin. However, this paxillin outline is not obvious in *siNFATC4*-transfected CCL-151 fibroblasts. The distribution of F-actin also shows a more relaxed cytoskeleton architecture in *siNFATC4*-transfected fibroblasts. In *siNC*-transfected cells, F-actin bundles are highly stretched on the non-adhesive edge of micropatterns (lower two rows, yellow arrows), which is the contraction edge, suggesting involvement in cell contraction. In contrast, these fibers are comparable more evenly distributed in *siNFATC4*-transfected fibroblasts. B. Quantification of MLT-018 nucleus area, circularity, and roundness. C. The shape and position of MLT-018 nuclei. Activated myosin was indicated by pMYL9 (gray), and the nucleus was indicated by DAPI (magenta). Statistics: *p < 0.05, **p < 0.01, ***p < 0.001, ****p < 0.0001, and n.s.: not significant according to the t-test.

### Supplementary Tables

**Table S1:**

Phospho quantification results from CCL-151 cells on substrates with different stiffness.

**Table S2:**

Table of significantly regulated phosphosites on different substrate rigidities (Anova 1 % FDR).

**Table S3:**

Phospho quantification results from the CCL-151 cells spreading time course.

**Table S4:**

Table of significantly regulated phosphosites from the spreading time course (Anova 1 % FDR).

**Table S5:**

Table with description of the function of proteins whose phosphorylation sites are regulated by substrate stiffness in the process of cell spreading in CCL-151 fibroblasts.

**Table S6:**

Table with description of the function of proteins whose phosphorylation sites are regulated by substrate stiffness in the cell line (CCL-151) as well as in primary cells (phLF).

**Table S7:**

Tables with information on used antibodies and primer pairs.

## Experimental procedures

### Cell culture

CCL-151 cell line was obtained from the American Type Culture Collection and tested negative for mycoplasma. Cells were maintained in Dulbecco’s Modified Eagle Medium/Nutrient Mixture F-12 (DMEM/F-12) (Thermo Fisher Scientific, #31330038) supplemented with heat-activated 10 % FBS (Sigma, #S0615-500ML) and 1 % penicillin/streptomycin (ThermoFisher Scientific, #15140122).

Primary human lung fibroblasts were obtained from the CPC-M bioArchive at the Comprehensive Pneumology Center (CPC Munich, Germany). The study was approved by the local ethics committee of the Ludwig-Maximilians University of Munich, Germany (Ethic vote #333-10). Written informed consent was obtained for all study participants. Human primary lung fibroblasts derived from peritumor control tissues and IPF-patient-derived lung explants were cultured in DMEM/F12 medium with 20 % FBS and 1% penicillin/streptomycin. All cells were cultivated at 37 °C and 5 % CO_2_ in a humidified atmosphere.

### Western blot, antibodies

For Western blot analysis, cells were lysed in RIPA buffer (50 mM Tris-HCl pH 7.6, 150 mM NaCl, 1 % NP-40, 0.1 % SDS, 0.5 % sodium deoxycholate) supplemented with complete protease inhibitor (Roche Diagnostics, Basel, Switzerland) and phosphatase inhibitor (PhosStop, Roche Diagnostics), incubated on ice for 30 min and centrifuged at 14,000 *g* for 10 min at 4 °C. Protein concentrations were determined using the Pierce^TM^ BCA Protein Assay Kit (ThermoFisher Scientific Inc., Waltham, MA, USA). 6x Laemmli loading buffer (6% SDS, 300 mM Tris-HCl, pH 6.8, 25 % (w/v) glycerol, 324 mM dithiothreitol (DTT), 0.1% bromophenol blue containing 50 mM NaF and 100 mM ß-glycerol-phosphate) was added to samples (10-20 µg protein lysate) and heated at 95 °C for 5 min. Lysates were loaded on 10-15% SDS-PAGE gels and blotted onto polyvinylidene difluoride (PVDF) membranes. Electrophoretic separation was performed at 100–120 V, RT and transfer to PVDF membranes was performed at 350 mA for 2 h on ice. Membranes were blocked in 1x Roti®Block (Roth, Karlsruhe, Germany) and incubated with primary antibodies o.n. at 4°C and subsequently with secondary antibodies 1 h, RT (1:20,000). ChemiDoc imaging system (BioRad Laboratories, Inc.) was used to detect chemiluminescence. Band intensities were quantified using Bio-Rad Image Lab software. Antibodies used were: Anti-GAPDH (1:1000, Cell Signaling #5174S), Anti-CK II alpha (1:1000, Abcam ab70774), Anti-Phospho-CK2 Substrate Motif [(pS/pT)DXE] (1:1000, Cell Signaling #8738), Anti-Phospho-Myosin Light Chain 2 (Thr18/Ser19) (1:2000, Cell Signaling #3674), Anti-Phospho-FoxO3a (S7) (1:1000, Cell Signaling 14724S), Anti-Phospho-Paxillin (S126) (1:1000, Life Technologies 441022G), Anti-Phospho-CBX3 (Ser93) (1:1000, Cell Signaling #2600), Anti-rabbit IgG, HRP-linked (1:20,000, Cell Signaling #7074) Anti-mouse IgG, HRP-linked (1:20,000, Cell Signaling #7074). Images were quantified in the Image Lab software (Biorad), calculated as the intensity of the band of interest divided by the intensity of the loading control for that lane.

### Phosphoproteome sample preparation

CytoSoft 6-well dishes (Advanced BioMatrix, Inc. San Diego, CA) with elastic modules 0.2, 0.5, 2, 8, 16 and 32 kPa or normal plastic dishes were coated with 10 µg/ml fibronectin (FN) for 1h, RT. Then, 500,000 cells were seeded per well for 120 min and lysed in 1 ml guanidinium chloride (GdmCl) lysis buffer (6 M GdmCl, 100 mM Tris pH 8.5, 10 mM Tris (2-carboxyethyl)phosphine (TCEP), 40 mM 2-chloroacetamide (CAA)).

Collected lysates were heated for 5 min, 95 °C. Lysates were cooled on ice for 15 min, sonicated with Bioruptor (Diagenode Inc., Denville, NJ) at maximum energy for 10 x 30 sec cycles and heated again for 5 min, 95 °C. Protein lysates were diluted 1:2 with water. Proteins were precipitated o.n. with 4 x volume of ice-cold acetone. Proteins were collected by centrifugation for 10 min at 4,000 *g* (4°C). Pellets were washed twice with ice-cold 80 % acetone, and air dried for 10 min, RT. Pellets were resuspended in 500 µL 2,2,2-trifluoroethanol (TFE) digestion buffer (10% TFE, 100 mM ammonium bicarbonate) with sonication in Bioruptor at maximum energy for 10 x 30 sec cycle or until a homogenous suspension was formed. Phosphopeptides were enriched according to the protocol as previously described (Humphrey *et al*, 2015)). Briefly, samples were digested overnight in 500 μl TFE digestion buffer. A first digestion was carried out with Lys-C 1:100 enzyme-to-protein ratio (wt/wt) at 37°C for 1h, 2,000 rpm, 37°C, followed by trypsin digestion o.n., 2,000 rpm, 37°C. 150 μl 3.2 M KCl, 55 μl 150 mM KH_2_PO_4_, 800 μl acetonitrile (ACN) and 95 μl trifluoroacetic acid (TFA) were added to the digested peptides. Peptides were mixed for 1 min, 2,000 rpm, RT, cleared by centrifugation and transferred to a clean 2 ml 96-well deep-well plate (DWP, Eppendorf, Hamburg, Germany). TiO_2_ beads (Titansphere^®^ Phos-TiO Bulk 10 µm, GL Science Inc., Tokyo, Japan, #5010-21315) were subsequently added to peptides at a ratio of 1:10 beads/protein, suspended in 80 % ACN/6% TFA, and incubated for 5 min, 2,000 rpm, 40 °C. Beads were pelleted by centrifugation for 1 min, 3,500 *g*, and the supernatant (containing non-phosphopeptides) was aspirated and discarded. Beads were suspended in wash buffer (60% ACN, 1% TFA) and transferred to a clean 2 ml DWP, and washed a further four times with 1 ml of wash buffer. After the final wash, beads were suspended in 100 μl transfer buffer (80 % ACN, 0.5% acetic acid), transferred onto the top of a C8 StageTip, and centrifuged for 3–5 min, 500 *g* or until no liquid remained on the StageTips. Bound phosphopeptides were eluted 2 times with 30 μl elution buffer (40 % ACN, 15 % NH_4_OH (25 %, HPLC grade)) each and collected by centrifugation into clean PCR tubes (Eppendorf). Samples were concentrated in a SpeedVac for 15 min, 45 °C and acidified with 10 μl 10 % TFA.

### StageTip desalting of peptides and phosphopeptides

Peptides were loaded onto styrenedivinylbenzene–reversed phase sulfonated (SDB-RPS; 3M Empore) StageTips in 100 μl SDB-RPS wash buffer (0.2 % TFA), and centrifuged for 3 min at 500 *g* or until ≤ 5 μl remained. StageTips were washed with a further 100 μl SDB-RPS wash buffer, and phosphopeptides were subsequently eluted with 60 μl SDB-RPS elution buffer (80 % ACN, 5 % NH_4_OH (25 %, HPLC grade)) into clean tubes by centrifugation for ∼5 min at 500 *g*. Samples were immediately concentrated in a SpeedVac for 30min at 45 °C, or until ∼2 μl remained. 7 μl MS loading buffer (2 % ACN, 0.3 % TFA) was added.

### Single-run MS/MS measurement

Peptides were loaded onto a 40cm column with a 75 μM inner diameter, packed in-house with 1.9 μM C18 ReproSil particles (Dr. Maisch GmbH). Column temperature was maintained at 50°C using a homemade column oven. An EASY-nLC 1000 system (Thermo Fisher Scientific) was connected to the mass spectrometer with a nanospray ion source, and peptides were separated with a binary buffer system of 0.1% formic acid (buffer A) and 60% ACN plus 0.1% formic (buffer B), at a flow rate of 300nl/min. Peptides were eluted with a gradient of 5–25% buffer B over 85 or 180min followed by 25–50% buffer B over 35 or 60min, resulting in ∼2- or 4h gradients respectively. Peptides were analyzed on a Q Exactive benchtop Orbitrap mass spectrometer (Thermo Fisher Scientific), with one full scan (300–1,600 *m*/*z*, *R* = 60,000 at 200 *m*/*z*) at a target of 3e^6^ ions, followed by up to five data-dependent MS/MS scans with higher-energy collisional dissociation (HCD) (target 1e5 ions, max ion fill time 120ms, isolation window 1.6m/z, normalized collision energy (NCE) 25%, underfill ratio 40%), detected in the Orbitrap (*R* = 15,000 at 200 *m*/*z*). Dynamic exclusion (40s or 60s) and apex trigger (4 to 7s) were enabled.

### LC-MS/MS analysis and data processing

MS raw files were processed using MaxQuant Version 1.5.3.34 (Tyanova *et al*, 2016). The spectra were searched with the Andromeda search engine using a false discovery rate (FDR) of < 0.01 on protein level as well as on peptide and modification level. In general the default settings of MaxQuant were used but with following changes. Methionine (M), acetylation (protein N-terminus) and phosphorylation (STY) were selected as variable modification. Carbamidomethyl (C) was selected as fixed modification. Furthermore, only peptides with a minimal length of seven amino acids were taken into account. The ‘match between runs’ option was turned on using a matching time window of 0.7 minutes. Peptide and protein identification was done by using the UniProt data base from human (April, 2018) which contained 20,138 entries. Each raw file was handled as one experiment. For the rigidity phosphoproteomic data set replicate 1 of 0.5 kPa was removed for the final analysis because of low identification number and outlier clustering within the replicates. Exactly the same was done with the phosphoproteomic spreading data set. Here the following replicates were not considered for the final analysis: replicate 2 of 20 min and replicate 4 of 30 min. Replicate 1 of 0.5 kPa, replicate was not considered for the final analysis due to low identification number and outlier clustering within the replicates.

### Bioinformatic and statistical analysis

#### Perseus

The processed data files were uploaded to the Perseus software (Tyanova *et al*, 2016) in order to perform bioinformatic data analysis. First, the label-free phosphorylation data were filtered to retain only these sites that have a localization probability > 0.75. In the next step, the data were rearranged, so that the different number of phosphorylations per peptide are not formatted as separate columns but just as one column per sample and three rows per phosphosite alternatively. The phosphorylation intensities were log2 transformed and filtered from reverse (decoy) database hits. The following protein annotations were added: GOBP name, GOMF name, GOCC name, GOPB slim name, GOMF slim name, GOCC slim name, KEGG name, Pfam, GSEA, Keywords and Corum. The different annotations were obtained from http://annotations.perseus-framework.org. Then, phosphosite-specific data such as sequence features, linear motifs, known sites, kinase substrate relations and regulatory sites were added to the data set obtained from the PhopshoSitePlus database. The respective replicates were grouped according to stiffness or spreading time.

Both datasets were filtered for valid values. In the rigidity data set, three values in at least one group had to be valid in order to pass the filter. For the spreading data set four values in at least one group had to be valid to not filter out the phosphorylation sites, since here five replicates per group were measured and not just four replicates as for the rigidity data set. The data were normalized by subtracting the median of each column.

Missing values were imputed using a width of 0.3 and a downshift of 1.8 for the total matrix. ANOVA testing for identification of regulated phosphosites was performed with and without prior imputation of the data. The phosphorylation sites found to be significantly regulated by ANOVA testing without imputation of the data were combined with the results of the performed ANOVA test with imputed values. In the next step, a matrix that only contained the significantly regulated phosphorylation sites was created. A principal component analysis was performed to further explore the data. Before proceeding with hierarchical clustering, the data was z-scored. This means, that the mean of each row is subtracted from each value and the result is dived by the standard deviation of the row. Based on the hierarchical clustering, the data were split into distinct clusters.

Relative enrichment of annotations for the IPA predicted CK2A1 regulated sites was achieved using Fisher’s exact test enrichment analysis (5% FDR). Ingenuity Pathway analysis. IPA is a web-based software application that facilitates the analysis, integration and understanding of data from gene expression, miRNA and SNP microarrays as well as metabolomics, proteomics and RNAseq experiments. IPA enables the search for specific information about genes, proteins, chemicals and drugs and the construction of interactive models of experimental systems. Data analysis and search functions help to understand the meaning of data, specific targets or candidate biomarkers in the context of larger biological or chemical systems. The software is supported by the Ingenuity Knowledge Base, which contains highly structured, detailed biological and chemical knowledge (IPA, Qiagen, Redwood City, www.qiagen.com/ingenuity).

The z-scored averages of the distinct significantly regulated phosphosites of each condition (stiffness or spreading time) were uploaded into the program. Observations were inferred and the identifier was assigned. After the data upload, a core phosphorylation analysis was performed using the z-scored intensity of the phosphosites as the base for analysis. Once all core analyses were calculated, a comparison analysis was run to obtain regulated Canonical Pathways, Upstream Regulators and Diseases and Functions. Canonical Pathways show the molecules of interest in known and well-established signaling pathways. Furthermore, if possible the likelihood is determined whether the pathway is activated or inhibited. Upstream analysis predicts which upstream regulators (any molecule that can affect the phosphorylation of another molecule) could be activated or inhibited to explain the phosphorylation changes in the data set. Diseases & Functions relates the molecules in the data set to known disease conditions and biological functions. Where applicable, IPA derives from the activity of the protein whether the associated disease or function is likely to be increased or decreased.

Metascape analysis. The members of the clusters defined by hierarchical clustering of the phospho data in Perseus were submitted to the web-based analysis tool Metascape (Zhou *et al*, 2019). This portal combines functional enrichment, gene annotation, membership search and interactome analysis of over 40 independent knowledge bases to comprehensively analyze OMICs-based data. After inserting the gene list, the express analysis was performed for each cluster. In the settings species h. sapiens was selected as input and also used for analysis. All pathway and process enrichment analysis for each cluster were combined in one heatmap.

Scanpy packages. Scanpy is a Python-based, scalable toolkit which was originally designed for the analysis of single-cell gene expression data (https://github.com/theislab/Scanpy) (Strunz *et al*, 2020; Wolf *et al*, 2018). Even though the analysis tool was intended for the analysis of single-cell gene expression data, it was used in this work for the analysis of phosphorylation intensity data. Scanpy can be used for clustering and visualization of data and can easily be run on Jupyter notebooks, which represents an interactive web-based environment (www.jupyter.org).

Tables generated in Perseus containing the significantly regulated phosphosites were exported and imported into Scanpy and a scanpy object was created. This was done for the CCL-151 stiffness dataset, the CCL-151 spreading dataset, the merged dataset spreading and stiffness and the combined phosphorylation data from CCL-151 and phLFs on different substrate rigidities. The merge of data sets was performed in Excel by making use of the unique identifiers for each phosphorylation site which were generated by processing all phospho data together in the proteomics software package MaxQuant.

### RNAi

Fibroblasts (Passages 7–10) were used in this study. Cells were dissociated with TrypLE Express Enzyme (1X), no phenol red-100 (Thermo Fisher Scientific, #12604013), and adjusted to a proper concentration using antibiotics-free culture medium before seeding in plates which precoated with 10 μg/mL FN1 (Sigma, #F1141-1MG). Synthetic Silencer Select siRNAs against human NFATC4 (Thermo Fisher Scientific, Assay ID-s9484 and s9483), siRNAs against YBX1 (Thermo Fisher Scientific, Assay ID-s9732 and s9733), and Silencer™ Select Negative Control No. 1 siRNA (Thermo Fisher Scientific, #4390843) were used for RNAi. Two siRNAs targeting the same gene were pooled as a mix. The Lipofectamine™ RNAiMAX (Invitrogen, #13778100) was used as the transfection reagent for delivering siRNAs into fibroblasts, and the transfection was according to the manufacturer’s instructions. Briefly, siRNAs and RNAiMAX were pre-diluted in Opti-MEM (Thermo Fisher Scientific, #31985062), and incubated for 5 min at room temperature. Subsequently, siRNA solution was added to the RNAiMAX solution and mixed gently. After 20–30 min of incubation at room temperature, the transfection complexes were added to the cell suspensions. In some experiments, cells were transfected again 48 h later and cultured for another 24 h before proceeding to the next step.

### Immunofluorescence staining and microscopy

Glass coverslips or ibidi microscopic dishes (0.5 kPa and 28 kPa) were coated with 10 µg/ml fibronectin (FN). Cells were seeded for 120 min. Then the cells were washed 3 times with PBS and fixed with 4 % PFA for 20 min at RT. Afterwards, cells were washed 3 times for 5 min with PBS/TBS and permeabilized with 0.1 % Triton X-100 in PBS/TBS for 10 min. After 3 times of washing for 5 min, cells were blocked with 5 % BSA in PBS for 30 min at RT and incubated with primary antibodies diluted in 1 % BSA in PBS/TBS o.n., 4 °C. After that, cells were washed 3 times for 5 min with PBS/TBS followed by 2 h incubation with the secondary antibodies diluted in 1 % BSA in PBS. After washing 3 times for 5 min with PBS/TBS, cells were incubated with DAPI for 10 min at RT, followed by washing with PBS/TBS 3 times for 5 min. Cells on glass coverslips were mounted with Fluorescent Mounting Medium, whereas cells on ibidi microscopic dishes were covered with PBS/TBS. Samples were examined using Zeiss LSM 710 or AXIO Imager. Antibody and dilution information was documented in the supplementary table “Table Antibodies and primers.xlsx”.

### ECM deposition

The ECM deposition assay was carried in 384-well CellCarrier plates (PerkinElmer, #6007550), and modified procedures were performed based on the previously described (Gerckens *et al*, 2021). Briefly, The plate was coated with 10 μg/mL FN1 (Sigma, # F1141-1MG) for 1 hour at RT before seeding cells. After the removal of FN1, 50 μL fibroblasts in the antibiotics-free culture medium were dispensed into a well at the density of 6,000 cells/well. Then, 5 μL of siRNA transfection mixture was added to cells, the plate was shaken gently, and incubated overnight. On the next day, the medium was removed and replaced with the starving medium (1% FBS), and after 24 h starvation, the new medium (1% FBS) with or without TGF-β (1 ng/mL) was changed. Three days later, the plate was used for ECM staining. For ECM staining, the medium was removed, and 20 μL of primary antibody mix in the starvation medium was added to each well. The plate was then put back into the cell culture incubator and incubated for 3 hours, subsequently added with 20 μL of the secondary antibody as well as Hoechst 33342 (1 μg/ml), and incubated for another 1 hour. The wells were briefly washed 3 times with DPBS and fixed by 4% PFA for 30 min at 37°C. The plate was subsequently washed once with PBS and supplemented with 60 μL of PBS in each well. Antibody and dilution information was documented in the supplementary table “**Table Antibodies and primers.xlsx**”.

### Collagen Gel Contraction Assay

Cell suspension (1.5 × 10^5^ cells/mL) was transferred to a 2 mL Eppendorf tube, and TGF-β was supplemented to the tubes (final concentration was 1.5 ng/mL) for TGF-β treated groups. Then, 0.5 volumes of Collagen I solution were added to the cell suspension. An appropriate volume (obtained from a prior titration) of 0.5M NaOH was then added to the mixture and mixed gently. The mixture of cell and collagen was immediately dispensed (500 μL/well) to a 24-well plate, and the plate was placed in a 37°C incubator for 20 min. The culture media with or without TGF-β (1 ng/mL) was added gently (600 μL/well) along the wall of the well. The gel pads were released from the well by gently running a 200 μL pipet tip along the gel edge. The plate was subsequently placed in a humidified incubator at 37°C, with 5% CO_2_. The plate was scanned with ChemiDoc Imaging System (Bio-Rad) at indicated time points (12 h, 24 h, 48 h, 72 h, 96 h, 120 h) to record the contraction of gels.

### Collagen I gel invasion assay

Collagen I matrix was prepared from rat tail Collagen I (ThermoFisher, A1048301), and modified procedures were performed based on the previously described (Oehrle *et al*, 2015). Briefly, 0.7 M NaOH was mixed with 1 M HEPES at 1:1 to make solution A, which was mixed with 20% FBS in 10× PBS at 1:1 to make solution B. Next, the collagen solution was prepared by gently mixing solution B and Collagen I at 1:4. The freshly prepared collagen solution was added to a 96-well plate (50 µL/well) carefully to avoid bubbles. The plate was then incubated at 37°C for 4 h for gel polymerization. Then, 2 × 10^4^ cells were seeded on top of the polymerized collagen gel in each well. Cells were allowed to invade the collagen matrix for 96 h in a 37°C, 5% CO_2_ incubator. After that, the wells were washed with 1× PBS, and the cells were fixed with 4% PFA for 30 min at RT, permeabilized with 0.25% Triton X-100 overnight, and stained with DAPI (1:1000), Alexa Fluor™ 647 Phalloidin (Thermo Fisher Scientific, #A22287) (1:150) and anti-αSMA antibody (1:250) overnight at 4°C (75 µL/ well). The wells were briefly washed with 1×PBS for 3 × 10 min and stained with Donkey anti-mouse-568 (1:500) for 1 h at 37°C. Antibodies were diluted in TBST containing 0.1% Tween. For imaging, Z-stack images of each well were acquired using an LSM 710 confocal microscope. For the collagen invasion assay, fifty images were collected at 500 µm distance. The 3D projections of the confocal images were generated using ImageJ. The confocal Z-stacks were also imported into Imaris software (Version 9.9.1, Oxford Instruments) for 3D visualization and nucleus number quantification. Each single nucleus was assigned a spot by the spot detection algorithm using similar parameters described previously (Burgstaller *et al*, 2013), and the spots were filtered according to their z-positions and counted using Imaris’ statistical analysis tool.

### Reverse transcription-quantitative polymerase chain reaction (RT-qPCR)

The mRNA expression abundance of genes in fibroblasts was assessed by RT-qPCR. Briefly, total RNA was isolated using Quick-RNA Microprept Kit (Zymo Research, #R1050), according to the manufacturer’s instructions. RNA concentrations were measured spectrophotometrically using Nanodrop 1000 (Thermo Scientific). After that, cDNA was synthesized from total RNA (100 ng) using GoScript™ Reverse Transcriptase System (Promega, #A5000), and the reverse transcription was 42°C for 1h. Quantitative PCR (qPCR) was then performed on LightCycler 480 Real-Time PCR System using the GoTaq qPCR Master Mix (Promega, #A6001), and the primers pairs are listed in Table S7. The cycling parameters for qPCR were as follows: initial activation at 95°C for 2 min, followed by 50 cycles of denaturation at 95°C for 5 s, annealing and extension at 60°C for 1 min. The 2^−ΔΔCt^ method was used to determine the relative abundance of mRNA, and *GAPDH* was used as endogenous control.

### Fibronectin micropattern experiments

Fibroblasts were re-transfected 48 h after the first transfection, and culture for another 24 h. For fibroblast dissociation, 50% TypLE (diluted with DPBS) was used and the DMEM/F12 medium containing 1% FBS was used to terminate the digestion. The fibroblasts were then centrifuged at 300 g for 10 min, and the cell pellet was suspended in 10mL of pre-warmed DEMD/F12 containing 1% FBS to wash once more time. Then, 25,000 Fibroblasts were suspended in 2 mL of pre-warmed medium and seeded in a 6-well plate placed with an L-shape micropatterned coverslip. The micropattern coverslip was pre-coated with 100 μL of 5 μg/mL FN for 1 h at 37°C.

The cells were allowed to attach to the coverslip for ∼30 min, with a brief checking of the attachment and spreading of cells under a microscope, cells can be washed with 2 mL of medium to remove floating cells. After another 2.5 h, cell spreading was checked to make sure that most of the cells were spread well on the large ‘L’-shaped patterns.

To fix cells, 2 mL of 4% PFA in PBS were added directly to the medium and incubated for 10 min at 37°C. Cells were washed with 2 mL of PBS and 0.2% Triton X-100 was used for permeabilization for 10 min at room temperature. After washing twice with 2 mL of PBS, cells were blocked with 100 μL of 2% BSA in PBS for 1 h at room temperature. After the removal of BSA, the slide was incubated with 150 μL of primary antibody (1:300 for anti-paxillin, BD Bioscience #610051; 1:150 for anti-pMLC, CST #3674S) at 37°C for 3 h. The slide was washed with PBS (3 x 5min). A drop (100 μL) of secondary antibody (1:1000)and Phalloidin (1:150, Invitrogen #A22287) was placed on a piece of Parafilm, and the coverslip was placed upside down on the antibody drop with its FN-coated side in contact with antibody. The coverslip was incubated with the antibody for 1 h and then 2 mL of DAPI (1 μg/mL in PBS) for 10 min at room temperature, washed with PBS, and finally mounted on a microscope slide with 25 μL of mounting medium.

The DAPI channel was split from fluorescent images and converted to its greyscale using ImageJ. The nuclei were outlined based on DAPI signals. The nucleus area, and roundness were calculated automatically with ImageJ (Rueden *et al*, 2017). Roundness was calculated according to 4 × area/(π × major axis^2^).

### Data availability

Proteome raw data and MaxQuant processing tables can be downloaded from the PRIDE repository (Perez-Riverol *et al*, 2019) under the accession number PXD039774.

## Acknowledgements

We gratefully acknowledge the provision of human biomaterial (primary human fibroblasts) and clinical data from the CPC-M bioArchive and its partners at the LMU Hospital, Asklepios Biobank Gauting and the Ludwig-Maximilians University of Munich. We thank the patients and their families for their support. We thank Anja Disovic and Marion Frankenberger from the CPC-M Bioarchive for excellent assistance. We thank Igor Paron and Christian Deiml for expert support of the proteomics pipeline. We thank Prof. Dr. Stefanie Eyerich and Dr. Manja Jargosch for support with fibroblast transfections. We also thank Anastasia van den Berg for excellent technical support. This work was supported by the Deutsche Forschungsgesellschaft (DFG; grant number SCHI 1378/1-1, ‘Functional proteomics of cellular mechanosensin’), the German Center for Lung Research (DZL), the Helmholtz Association, and the Max Planck Society.

## Author contributions

HBS conceived the research narrative and experimental design and supervised the project. HBS and LFM wrote the paper. LFM and HBS performed proteomics experiments and analyzed the data. ZZ performed NFATC4 loss of function experiments and analyzed the data. ZZ, LFM, XW, SA, and AAW performed validation experiments. LFM, HBS, CHM and MA performed bioinformatic data analysis of phosphoproteomic data. NK, MGS, and JB provided patient material and resources. JP generated adhesive micropatterns. AÖY, GB, MM and HBS provided resources and funding. All authors read and approved the manuscript.

