## Supplementary material for "Phosphoproteomics of cellular mechanosensing reveals NFATC4 as a regulator of myofibroblast activity": suplementary tables: Table S5_proteins-of-phosphosides-regulated-by-spreading-and-stiffness.docx

**Supplementary table SX**

Description of the function of proteins whose phosphorylation sites are regulated by **substrate stiffness** **and cell spreading**. The provided protein information was taken from the UniProt database ([www.uniprot.org](http://www.uniprot.org)).

| Protein | Phosphosite | Cluster | Description | Uniprot |
| --- | --- | --- | --- | --- |
| ABCC1 | T918 | A | Mediator of export of drugs and organic anions from the cytoplasm | P33527 |
| ABCF1 | S105, 108, 109 | A | Required for mRNA translation initiation | Q8NE71 |
| ARHGEF2 | S174, 177 | A | Activator of Rho-GTPases involved in many cellular processes such as cell motility and polarization | Q92974 |
| ATRX | S875, 876 | A | Transcriptional regulator which is also involved in chromatin remodeling | P46100 |
| CAMK2D | T337 | A | Calcium/calmodulin dependent protein kinase implicated in the control of Ca2+ homeostasis and excitation-contraction coupling | Q13557 |
| CEP170 | S1169, 1174 | A | Centrosomal protein involved in microtubule organization | Q5SW79 |
| EHBP1 | S428 | A | Probably involved in actin reorganization and links clathrin-mediated endocytosis to the cytoskeleton | Q8NDI1 |
| GPATCH2 | S115, 117 | A | G patch domain containing protein 2-like | Q9NWQ4 |
| GRK5 | S484, T485 | A | Serine/threonine kinases which phosphorylates mainly activated GPCRs | P34947 |
| HIRIP3 | S289, 291 | A | Interacts with HIRA and histones and thus might be important for chromatin function and histone metabolism | Q9BW71 |
| HNRNPUL2 | S169, T165 | A | Protein involved in RNA binding | Q1KMD3 |
| MAP1A | S2237 | A | Structural protein participating in the filamentous bridging between microtubules and other cytoskeletal elements | P78559 |
| MECP2 | S357 | A | Promotes transcriptional repression through interaction with histone deacetylase and the corepressor SIN3A | P51608 |
| MICAL | S872, 875 | A | Nuclear monooxygenase that enhances the depolymerization of F-actin by facilitating the oxidation of specific methionine residues on actin to generate methionine sulfoxide, which leads to the disintegration of actin filaments and prevents repolymerization | O94851 |
| MYO18A | S2041, 2043 | A | Might connect the Golgi membranes to the cytoskeleton and contribute in the tensile force required for the vesicles to sprout from the Golgi | Q92614 |
| NDRG1 | T328 | A | Tumor suppressor in various cell types and regulator of microtubule dynamics | Q92597 |
| SF3B1 | S332, T326, 328 | A | Component of the SF3B splicing factor complex and thus important in pre-mRNA splicing | O75533 |
| SPECC1L | S832 | A | Cytospin-A is essential in actin organization and microtubule stabilization and therefore involved in cell adhesion and migration | Q69YQ0 |
| SRRM1 | S389, 391, 393 | A | Contributing to numerous processes in pre-mRNA processing also as part of multiprotein mRNP complexes | Q8IYB3 |
| SRRM2 | S778, 780, 783, 1497, 1499 | A | Component of the spliceosome | Q9UQ35 |
| SRSF2 | S206, 208, 212 | A | The factor is needed for formation of the ATP-dependent splicing complex and the pre-mRNA splicing | Q01130 |
| AFAP1 | S664 | B | Actin filament-associated protein 1, crosslinker of actin filaments to bundles and networks, adapter molecule which links proteins to actin | Q8N556 |
| AKAP12 | S696, 698 | B | Anchoring protein which regulates the compartmentation of PKA and PKC | Q02952 |
| APC | S2242, 2244 | B | Stabilizer of microtubules which regulates actin fiber dynamics | O95996 |
| AHRGAP21 | S1712 | B | The depletion of ARHGAP21 initiates cell proliferation and accumulation of F-actin stress fibers | Q5T5U3 |
| C2orf49 | S189, 193 | B | Protein of the ashwin family | Q9BVC5 |
| CBX3 | S97 | B | Chromobox protein homolog which is probably part of heterochromatin-like complexes involved in transcriptional silencing | Q13185 |
| CEP170 | S452 | B | Centrosomal protein involved in microtubule organization | Q5SW79 |
| CTNND1 | S861 | B | Catenin delta-1 is a key regulator of cell-cell adhesion and is also involved in gene transcription. | O60716 |
| DPYSL2 | S522 | B | Involved in neuronal developement, neuronal growth, cell migration and semaphorin class 3 signaling with subsequent cytoskeletal rearrangement | Q16555 |
| FAM129B | S641 | B | Protein Niban 2 might play a role in apoptosis suppression | Q96TA1 |
| FN1 | S2475 | B | Fibronectin binds ECM proteins and cell surfaces and is involved cell adhesion and motility as well as cell shape maintenance | P02751 |
| GOLPH3 | S35, 36 | B | Golgi phosphoprotein 3 links the cytokeleton with the Golgi membranes and is involved in vesicle budding from the Golgi | Q9H4A6 |
| GTF3C1 | S1062, 1068 | B | General transcription factor 3C polypeptide 1 is needed for transcription mediated by RNA polymerase III | Q12789 |
| HDGFRP2 | S232, 234 | B | Regulates expression of Cyclin D1 therby controlling cellular growth | Q7Z4V5 |
| HDLBP | S944 | B | Vigilin may play a role in protection from over-accumulation of cholesterol | Q00341 |
| HNRNPC | S241 | B | The protein may be involved in pre-mRNA splicing and the start of spliceosome assembly | P07910 |
| HNRNPUL2 | S185 | B | Protein involved in RNA binding | Q1KMD3 |
| HTATSF1 | S713, 714 | B | General transcription factor important for transcriptional elongation | O43719 |
| IWS1 | S511, 513 | B | Transcription factor which defines the composition of RNA polymerase II elongation complex | Q96ST2 |
| KIAA1462 | S755, 757 | B | Involved in cell adhesion | Q9P266 |
| LEO1 | T629, S630 | B | Component of the PAF1 complex which is involved in transcription of Hox and Wnt target genes | Q8WVC0 |
| LMNA | S404, 407, 414 | B | Component of the framework for the nuclear envelope, the so called nuclear lamina. Important for chromatin organization and nuclear assembly | P02545 |
| MEPCE | T213 | B | Part of 7SK RNP complex and has an enzymatic methyltransferase activity | Q7L2J0 |
| MSL3 | S400, T405 | B | Involved in transcriptional regulation and chromatin remodeling | Q8N5Y2 |
| NAB2 | S162, 171 | B | Transcriptional repressor for the zinc finger transcription factor EGR1 and EGR2 | Q15742 |
| PCIF1 | T150 | B | Cap-specific adenosine methyltransferase | Q9H4Z3 |
| PRKAA1 | T490 | B | Catalytic subunit of AMP-activated protein kinase (AMPK), an energy sensor protein kinase that plays a crucial role in controlling cellular energy metabolism | Q13131 |
| PVR | S406 | B | Promotes NK cell adhesion and enables NK cell effector functions | P15151 |
| S1PR2 | S332 | B | Receptor for the lysosphingolipid sphingosine 1-phosphate | O95136 |
| SNX1 | T41 | B | Engaged many stages of intracellular trafficking | Q13596 |
| SRRM1 | S207, 209, 211, 308 | B | Contributing to numerous processes in pre-mRNA processing also as part of multiprotein mRNP complexes | Q8IYB3 |
| SRRM2 | S248, 250, 1218, 1219, 1582 | B | Component of the spliceosome | Q9UQ35 |
| STK11IP | S493 | B | Controls function of kinase STK11/LKB1 via the subcellular localization | Q8N1F8 |
| TJP1 | S178 | B | Scaffold protein which connect tight junction transmembrane proteins for example claudine, compound adhesion molecules and occludin to actin | Q07157 |
| TOP2B | S1550, 1552 | B | DNA topoisomerase inducing double-strand breaks | Q02880 |
| UBAP2L | S116 | B | Has an important role in the activation of the long-term repopulation of haematopoietic stem cells | Q14157 |
| VAMP4 | S88, 90 | B | The vesicle-associated membrane protein 4 might be a marker for the sorting pathway which is important for the remodeling the secretory response of granule | O75379 |
| ZMYND8 | S475 | B | Possible transcriptional corepressor for KDM5D and involved in down-regulation of many metastasis-associated genes | Q9ULU4 |
| ACIN1 | S655, 657 | C | Part of the splicing-dependent multiprotein exon junction complex which is located at splice junctions on mRNAs | Q9UKV3 |
| -- | S73, 79 | C |  | B4DZQ5 |
| CCNL1 | S335 | C | Cyclin-L1 plays a role in pre-mRNA splicing | Q9UK58 |
| CEP170B | S421 | C | Involved in microtubule organization | Q9Y4F5 |
| CILP2 | S207 | C | The protein might be involved in cartilage scaffolding | Q8IUL8 |
| DBN1 | S142 | C | Drebrin is an actin-cytoskeleton-organizing protein and is important in cell projections | Q16643 |
| DTD1 | S196 | C | ATPase probably important in DNA replication | Q8TEA8 |
| FAM129B | S641, 646 | C | Protein Niban 2 might play a role in apoptosis suppression | Q96TA1 |
| FOXK1 | S213, 223 | C | Forkhead box protein K1 which can be a transcriptional activator or repressor depending on context playing a role in various processes for example glucose metabolism | P85037 |
| HIRIP3 | S196, 199 | C | Interacts with HIRA and histones and thus might be important for chromatin function and histone metabolism | Q9BW71 |
| IWS1 | S248, 250, 398, 400 | C | Transcription factor which defines the composition of RNA polymerase II elongation complex | Q96ST2 |
| KIAA1462 | S757 | C | Involved in cell adhesion | Q9P266 |
| LARP7 | S258, 261 | C | Binding protein of small nuclear RNA which regulates their function and processing | Q4G0J3 |
| LMNA | S12, 22 | C | Component of the framework for the nuclear envelope, the so called nuclear lamina. Important for chromatin organization and nuclear assembly | P02545 |
| MAP1A | S612, T616 | C | Structural protein participating in the filamentous bridging between microtubules and other cytoskeletal elements | P78559 |
| MAP1B | S992, 995 | C | Involved in microtubule polymerization and stabilization | P46821 |
| MAP4 | S280 | C | A non-neuronal microtubule-associated protein which enhances microtubule assembly | P27816 |
| MAPK3 | T202, Y204 | C | Serine/threonine kinase with an key role in MAP kinase signal transduction. Various cellular functions such as cell growth and survival, adhesion and differentiation are regulated by regulation of transcription and cytoskeletal rearrangements | P27361 |
| MARCKS | S135, T143 | C | MARCKS is a filamentous (F) actin cross-linking protein and the most prominent cellular substrate for protein kinase C and binds the proteins calmodulin, actin and synapsin | P29966 |
| MYLK | S1773, 1779 | C | Calcium/calmodulin-dependent myosin light-chain kinase involved in smooth muscle contraction through phosphorylation of myosin light chains (MLC). In addition, it controls the actin-myosin interaction through a non-kinase activity. | Q15746 |
| NES | S680 | C | Nestin enhances the degradation of phosphorylated vimentin intermediate filaments (IF) during mitosis and could be implicated in the trafficking of IF proteins | P48681 |
| PCYT1A | S319, 323 | C | The protein regulates phosphatidylcholine synthesis | P49585 |
| PHIP | S1281, 1283 | C | Involved in the control of cell morphology and cytoskeletal organization. | Q8WWQ0 |
| PHRF1 | S1359, 1360 | C | Protein containing a PHD and RING finger domain | Q9P1Y6 |
| PTPN14 | S465 | C | Protein tyrosine phosphatase, which is important in the regulation of lymphangiogenesis, cell-cell adhesion, cell-matrix adhesion, cell migration and growth and also regulates TGF-beta gene expression | Q15678 |
| PXN | S258, 261 | C | Cytoskeleton protein implicated in actin membrane fixation in focal adhesions | P49023 |
| RRBP1 | S713 | C | The protein functions as a ribosome receptor and facilitates the interaction between the ribosome and the endoplasmic reticulum membrane | Q9P2E9 |
| SART3 | S10 | C | U6 snRNP-binding protein that acts as a recycling factor of the splicing machine | Q15020 |
| SLC38A1 | S52 | C | Acts as a sodium-dependent amino acid transporter | Q9H2H9 |
| SLC39A7 | S275, 276 | C | Zinc transporter, which transports Zn2+ from the endoplasmic reticulum/Golgi apparatus to the cytosol | Q92504 |
| SRRM2 | S778, 780, 952, 954, 1219 | C | Component of the spliceosome | Q9UQ35 |
| STX12 | S139, T142 | C | SNARE protein which regulates protein transport between late endosomes and the trans-Golgi network | Q86Y82 |
| TBX2 | S653, 657 | C | T-box transcription factor which regulates genes required for mesoderm differentiation | Q13207 |
| WIZ | S1017 | C | Might link the histone methyltransferases EHMT1 and EHMT2 in a heterodimer and stabilizes the complex | O95785 |
| YBX1 | S174, S176 | C | DNA and RNA binding protein implicated in many processes like RNA stabilization, splicing and transcription regulation | P67809 |
