## Supplementary material for "Phosphoproteomics of cellular mechanosensing reveals NFATC4 as a regulator of myofibroblast activity": suplementary tables: Table S6_proteins-of-phosphosides-regulated-by-stiffness-inCCL151-and-phLF.docx

**Supplementary table SX**

Description of the function of proteins whose phosphorylation sites are regulated by **substrate stiffness** **in the cell line (CCL151) as well as in primary cells (phLF)**. The provided protein information was taken from the UniProt database ([www.uniprot.org](http://www.uniprot.org)).

| Protein | Phosphosite | Louvain | Description | Uniprot |
| --- | --- | --- | --- | --- |
| AAK1 | S618, T620 | 1 | AP2-asscoated protein kinase 1 involved in clathrin-mediated endocytosis and controls phosphorylation of other AP2 subunits and their cellular localization | Q2M2I8 |
| ABLIM3 | S503, 504 | 1 | Actin-binding LIM protein 3 is probably a scaffold protein and regulates ABRA activity as well transcriptional activation of ABRA-dependent SRF | O94929 |
| AHNAK | S5749, 5752 | 1 | Might be needed for neuronal cell differentiation | Q09666 |
| ATXN2L | S594 | 1 | Contributes to the regulation of the development of stress grains and P-bodies | Q8WWM7 |
| BCL9L | T957 | 1 | Transcriptional activator which enhances ß-catenin transcription and is important in development of tumors | Q86UU0 |
| BOD1L1 | S484, 484 | 1 | Part of the fork protection machine necessary to protect stalled/damaged replication forks from uncontrolled DNA2-dependent resection. | Q8NFC6 |
| CEP170 | S802 | 1 | Centrosomal protein involved in microtubule organization | Q5SW79 |
| CEP170B | S1545, 1548 | 1 | Involved in microtubule organization | Q9Y4F5 |
| CHAF1B | S429, T433 | 1 | Chromatin assembly factor part of a complex which may control chromatin assembly in DNA repair and replication | Q13112 |
| CHAMP1 | S204, 214 | 1 | Necessary for the correct alignment of chromosomes at the metaphase and their accurate separation during mitosis; Contributes to maintaining the binding of the spindle microtubules to the kinetochore during sister chromatid biorientation | Q96JM3 |
| CNN3 | S323 | 1 | Thin filament-associated protein involved in the regulation and modulation of smooth muscle contraction. It is able to bind to actin, calmodulin, troponin C and tropomyosin. | Q15417 |
| DBN1 | S339, 345 | 1 | Drebrin is an actin-cytoskeleton-organizing protein and is important in cell projections | Q16643 |
| ERCC5 | S562, 563 | 1 | Single-stranded structure-specific DNA endonuclease important in DNA excision repair | P28715 |
| KAT7 | S124, T128 | 1 | Histone acetyltransferase which is part of HBO1 complexes responsible for acetylation of histone H3 at Lys14 thereby mediating processes such as transcription, immune regulation and ubiquitination | O95251 |
| MAP1A | S1326, 1329 | 1 | Structural protein participating in the filamentous bridging between microtubules and other cytoskeletal elements | P78559 |
| MAP1B | S831, 1256 | 1 | Involved in microtubule polymerization and stabilization | P46821 |
| MIEF1 | S59 | 1 | Protein of the mitochondrial outer membrane responsible for regulation of mitochondrial fission | Q9NQG6 |
| MYH10 | S1975 | 1 | Cellular myosin important in cytoskeleton reorganization, focal adhesion formation, and lamellipodial extension; The function is mechanically antagonized by MYH9 | P35580 |
| MYL9 | S20 | 1 | Regulatory subunit of myosin, which is involved in regulating the contractile activity of both smooth muscle and non-muscular cells via phosphorylation; Important in cytokinesis, receptor capping and cell movement | P24844 |
| MYO9B | S114, 115 | 1 | Unconventional myosins are needed for intracellular movements. Binding actin with high affinity in the absence and presence of ATP, its mechanochemical activity is blocked by calcium ions | Q13459 |
| NCOA7 | S208, 2011 | 1 | Coactivator mediating the transcriptional activity of various nuclear receptors | Q8NI08 |
| NEXN | S160, 162 | 1 | Nexilin plays an important role in cell migration via association with actin | Q0ZGT2 |
| NMD3 | S468, T470 | 1 | Ribosomal export protein which functions as an adaptor for export of the 60S ribosomal subunit | Q96D46 |
| NUP188 | S1709 | 1 | Component of the nuclear core complex | Q5SRE5 |
| NUP214 | S430 | 1 | Component of the nuclear core complex and involved in nucleocytoplasmic transport | P35658 |
| PHF14 | S298, 302 | 1 | Protein with zinc finger binding motif | O94880 |
| PHIP | S674 | 1 | Involved in the control of cell morphology and cytoskeletal organization | Q8WWQ0 |
| PPIP5K2 | S1016 | 1 | Bifunctional inositol kinase that functions together with the IP6K kinases IP6K1, IP6K2 and IP6K3 | O43314 |
| PTPN14 | S620 | 1 | Protein tyrosine phosphatase, which is important in the regulation of lymphangiogenesis, cell-cell adhesion, cell-matrix adhesion, cell migration and growth and also regulates TGF-beta gene expression | Q15678 |
| PTRF | S167 | 1 | Important for caveolae formation and organization in all tissues | Q6NZI2 |
| RPS6KB1 | T444, S447 | 1 | Serine/threonine protein kinase that acts downstream of mTOR signalling in a reaction to growth factors and nutrients to stimulate cell proliferation, cell growth and cell cycle progression | P23443 |
| RTKN | S220 | 1 | Rhotekin promotes the Rho signal to enable NF-kappa-B activation and could lead to increased resistance to apoptosis | Q9BST9 |
| SF3B1 | T326, 328 | 1 | Component of the SF3B splicing factor complex and thus important in pre-mRNA splicing | O75533 |
| SIPA1 | S55 | 1 | Activator of the GTPase for nuclear Ras-related regulatory proteins Rap1 and Rap2 | Q96FS4 |
| SMTN | S792 | 1 | Smoothelin is a cytoskeletal structure protein | P53814 |
| TCOF1 | S171, T173 | 1 | Nucleolar protein which functions as a regulator of RNA polymerase I | Q13428 |
| THUMPD1 | S86, 88 | 1 | Works as a tRNA binding adapter to facilitate NAT10 dependent tRNA acetylation | Q9NXG2 |
| TMF1 | S338, 344 | 1 | TATA element modulatory factor is a potential coactivator of the androgen receptor and modulates STAT3 degradation. | P82094 |
| TNS1 | S1446, 1454 | 1 | Tensin-1 is implicated in the formation of fibrillar adhesions and May be implicated in cell migration, cartilage development and in the interconnection of signal transduction pathways with the cytoskeleton | Q9HBL0 |
| TRIM16 | T55 | 1 | E3 ubiquitin ligase, which has an essential role in the regulation of autophagic response and ubiquitination in lysosomal and phagosomal defects. | O95361 |
| ULK1 | S450 | 1 | Serine/threonine protein kinase which is implicated in autophagy in reaction to starvation. | O75385 |
| BRD1 | S1183, 1186 | 4 | Bromodomain-containing protein 1 is a scaffold subunit of many histone acetyltransferase complexes | O95696 |
| CAMK2D | T337 | 4 | Calcium/calmodulin dependent protein kinase implicated in the control of Ca2+ homeostasis and excitation-contraction coupling | Q13557 |
| EHBP1 | S434, 436 | 4 | Might contribute to actin reorganization and links clathrin-mediated endocytosis to the actin cytoskeleton. | Q8NDI1 |
| FLNA | S2152, 2158 | 4 | Filamin-A stimulates orthogonal branching of actin filaments and connects actin filaments with membrane glycoproteins. In addition, the protein anchors a variety of transmembrane proteins to the actin cytoskeleton and provides a scaffold for a vast range of cytoplasmic signalling proteins. | P21333 |
| IWS1 | S248, 398, 400 | 4 | Transcription factor which defines the composition of RNA polymerase II elongation complex | Q96ST2 |
| MARCKS | S135, T143 | 4 | MARCKS is a filamentous (F) actin cross-linking protein and the most prominent cellular substrate for protein kinase C and binds the proteins calmodulin, actin and synapsin | P29966 |
| MYL9 | T18, S19 | 4 | Regulatory subunit of myosin, which is involved in regulating the contractile activity of both smooth muscle and non-muscular cells via phosphorylation. Important in cytokinesis, receptor capping and cell movement | P24844 |
| MYLK | S1773, 1776, 1779 | 4 | Calcium/calmodulin-dependent myosin light-chain kinase involved in smooth muscle contraction through phosphorylation of myosin light chains (MLC). In addition, it controls the actin-myosin interaction through a non-kinase activity. | Q15746 |
| RAI14 | S293 | 4 | Ankycorbin controls actin regulation in ectoplasmic specialization, a type of testicular cell junction. | Q9P0K7 |
| RTN4 | S15 | 4 | Reticulon-4 is involved in induction of and stabilization of endoplasmic reticulum tubules | Q9NQC3 |
| SRRM1 | S289, 391, 393 | 4 | Contributing to numerous processes in pre-mRNA processing also as part of multiprotein mRNP complexes | Q8IYB3 |
| SRRM2 | S952, 954 | 4 | Component of the spliceosome | Q9UQ35 |
| TRIP10 | S296 | 4 | Facilitates CDC42-induced actin polymerization by recruitment of WASL/N-WASP, which then activates the Arp2/3 complex | Q15642 |
